## Supplementary Material for "Tracing cancer evolution and heterogeneity using Hi-C"

### Contents

**1 Notation & Setup** **2**

**2 Covariate correction** **3**

**3 Inference of copy numbers & mixture proportions** **4**

**4 Inference of large-scale structural variants** **10**

**5 Proof of Theorem** **14**

**6 Supplementary Figures** **17**

### Supplementary Note 1

HiDENSEC broadly proceeds by (i) correcting Hi-C counts for confounding covariates, (ii) using these corrected counts to infer mixture proportion and copy number, and (iii) identifying large structural variants based on the resulting profiles. Before describing each of these steps in detail, and in order to make equations easier to understand, we begin by fixing the notation and introducing general conventions that will be used throughout.

### 1 Notation & Setup

Our starting point is a raw nucleotide-level Hi-C matrix $H^{\text{raw}}$ obtained by mapping Hi-C or Fix-C reads to a reference genome (**Online Methods**). This matrix is symmetric, i.e., $H^{\text{raw}}_{ij} = H^{\text{raw}}_{ji}$.

In order to balance signal-to-noise ratios and computational burden with interpretability and localization of changes in copy number and/or rearrangement breakpoints/junctions, we first coarse grain this nucleotide-level Hi-C matrix $H^{\text{raw}}$ to genomic windows of length $w = 50\text{kb}$:

$$H_{ij} = \sum_{\substack{m=i,\ldots,i+w \\ n=j,\ldots,j+w}} H^{\text{raw}}_{mn},$$

The indices of this matrix label genomic windows. All further steps in HiDENSEC work with such a coarse-grained matrix.

For a completely homogeneous cell population, we can model $H$ as a random matrix of the form $H = Nh + \varepsilon_N$, where $N$ is the total number of cells involved in the Hi-C experiment, $h$ represents the (deterministic) interactions between genomic loci in cells of a given type (and so in particular, $h$ may differ across different cell populations), and $\varepsilon_N$ is an error matrix, on which we do not impose any restrictions other than $\lim_{N\to\infty} \varepsilon_N/N = 0$ (in probability). This assumption ensures the identifiability of $h$, and is expected under the very plausible assumption that the numbers of reads contributed by different cells in the experiment are independent).

Hi-C experiments are generally conducted on samples containing mixtures of cells with different genomes. We denote the set of genomes by $\mathcal{G}$, and represent the cellular mixture fractions by $\left(f^G\right)_{G\in\mathcal{G}}$. Since chromatin contacts only occur between genomic segments within the same nucleus, the Hi-C contact map of the sample is expected to be a simple superposition of these distinct Hi-C signatures. That is,

$$h_{ij} = \sum_{G\in\mathcal{G}} f^G h^G_{ij},$$

where $h^G$ is the $h$ matrix associated with genome $G$. To test the superposition assumption we confirmed the lack of Hi-C contacts between neighboring cells by performing Fix-C on a 1:1 mixture of human and mouse cells, finding negligible ligation products that include both mouse and human sequence (**Supplementary Table 1**).

Each genome $G$ will have an absolute copy number profile $p^G_i$ at each site $i$ representing the local, integer-valued, copy number at that locus. In a superposition of multiple genomes, however, we more directly measure the weighted average of these copy numbers, $\pi_i = \sum_{G\in\mathcal{G}} f^G p^G_i$, which we refer to as the effective copy number $\pi_i$ for the sample. If we knew the contact maps $h^G_{ij}$ in isolation for each genome $G \in \mathcal{G}$, then we should be able to infer the mixture proportions $f^G$, for $G \in \mathcal{G}$. With these in hand, we can also interpret rearrangements within each genome, which are represented by $p_{ij}^G$, the absolute number of site $i$ copies that are in contact with site $j$ in genome $G$ due to translocations and other rearrangements. To keep formulas and descriptions compact, we will occasionally refer to $p_i$ as $p_{ii}$.

In analyzing the Hi-C matrices discussed in this paper, we find that it is typically sufficient to include one or sometimes two cancer genotypes along with the diploid reference genome (which is free of copy number and other structural variation). Thus $|\mathcal{G}| = 2$ for most samples, except sample 1 from patient 3 which requires $|\mathcal{G}| = 3$. When $|\mathcal{G}| = 2$ there is no risk of confusing distinct genomes, and so for notational simplicity we drop the superscripts and refer to the interaction strengths, mixture proportion and copy numbers of the non-reference genome as $h$, $f$ and $p$, respectively.

The goal of HiDENSEC is then simply stated: Given experimentally observed Hi-C counts $H$ from a composite sample, we wish to infer the mixture proportions $f$, copy number profile $p = (p_i)$, and identify the set of non-adjacent structural variants $\{(i, j) : p_{ij} > 0\}$ together with the precise nature of these structural changes; that is, determine the values of $p_{ij}$ (whenever identifiable).

### 2 Covariate correction

As described in the **Online Methods**, there are several biological and technical effects which can substantially affect read counts (see **Supplementary Figure 1A**, and **Supplementary Figure 1B**). To begin to account for these covariates we focus on the "diagonal" local contact $h_{ii}$ and model its dependence on the true contacts $p_i$ as

$$h_{ii} = C_0 p_i g(x_i^0, x_i^1, x_i^2, x_i^3), \tag{1}$$

for a function $g$ to be determined, where $x^0$ through $x^2$ encode the numerical, experiment-related covariates (e.g., GC content, cut-site density, and read mappability), and $x^3$ is a discrete indicator of one of the six chromatin compartments $\mathcal{C} = \{\text{A0, A1, A2, B1, B2, B3}\}$.

Two observations inform the form of $g$:

1. The dependence between Hi-C contact intensities and covariates appears to be sensitive to protocol differences in the experiment. Although *in-situ* Hi-C based covariate structures tend to look very similar to **Supplementary Figure 1**, those obtained from Fix-C experiments usually look like the plots depicted in **Supplementary Figure 2**.

2. Even though the details of this effect may differ across protocols, their qualitative shape appears close to linear for $x^0$ and $x^1$, with $x^2$ exhibiting a cut-off phenomenon (see panels B of **Supplementary Figure 1** and **Supplementary Figure 2**) without substantial interaction (see panels C of the same figures). Constructing $g$ using these qualitative trends leads to satisfactory model fit (see the subsequent paragraph).

We therefore filter genomic bins, retaining only those for which $x^2 > 0.8$, and adopt a simple descriptive linear regression model, with different regression line for each type of compartment for $x^0$ and $x^1$. That is, $g$ is fit as

$$g(x^0, x^1, x^2, x^3) = \sum_{c \in \mathcal{C}} \mathbb{1}_c(x^3) g_c(x^0, x^1), \tag{2}$$

where $g_c$ is linear for all $c \in \mathcal{C}$. For purely diploid genomes this model explains 80-90% of the observed variance, satisfying standard model-fit criteria (normality and identical distribution of studentized residuals, independence of residuals and predictions, etc.)

For simplicity we also assume that covariate corrections Eq. (1) and Eq. (2) derived for karyotypically normal genomes can also be applied to cancer genome (that is, as a first approximation we neglect changes in compartment structure in the cancer genomes). By the superposition principle it follows that Eq. (1) and Eq. (2) also hold with $p_i$ replaced by $\pi_i$. Such covariate correction was performed for all Hi-C maps used in the main manuscript, based on $g$ fit to Fix-C from a karyotypically normal reference, corresponding to Sample 1 - I. An illustration of both the need for adjusting raw counts, as well as the protocol sensitivity of any such adjustment is shown in **Supplementary Figure 11**.

In principle, one could attempt a similar correction procedure on the off-diagonal entries $h_{ij}, i \neq j$, with a corrector function $g' : (\mathcal{C} \times \mathbb{R}^3)^2 \to \mathbb{R}$ that depends on pairs of covariates. However, as indicated in the main text, the magnitude of off-diagonal signals are difficult to interpret due to uncertainty in the precise break-point within a 50 kb window, and the possibility of substantially altered compartment structure associated with large-scale structural variants (**Supplementary Figure 4**). Nevertheless, these off-diagonal signals can clearly distinguish $h_{ij} > 0$ from $h_{ij} = 0$. For the precise inference of large-scale structural variants and their copy numbers, this type of on-or-off signal is sufficient as will be illustrated in Section 4.

**Remark.** *If a reference contact map is not available, or the underlying experimental protocol is unknown, then an internal covariate correction is still possible based on read counts reliably identified to correspond to $\pi_{mode}$ (see section below). Such correction empirically performs competitively with the procedure described above, and in HiDENSEC is resorted to automatically if no explicit reference protocol and/or data set is specified.*

### 3 Inference of copy numbers & mixture proportions

Given the covariate correction as described in Section 2 the resulting corrected read counts $H_{ii}$ are effectively modeled as $H_{ii} = C_0 N \pi_i + \varepsilon_{ii}$ as mentioned in Section 1 (see also **Supplementary Figure 12A** for an illustration of the underlying generative model).

First we note that, in the general case (arbitrary numbers and forms of cancer genomes), we cannot infer $\pi$ from $H$ as a matter of principle, even in the limit of infinite data, without further assumptions:

1. *Relative copy number profiles determine absolute copy number profiles only up to integer scaling.* Since $\pi = \sum_{g \in \mathcal{G}} f^G p^G$ is only observed up to an overall factor, any two sets of copy number vectors $p$ and $p'$ that differ by a multiplicative constant (that is, $p' = Kp$ for some $K \in \mathbb{N}$) are indistinguishable on the level of $H$, since $C_0$ and $N$ are generally unknown. E.g., a completely diploid and a completely triploid genome are indistinguishable based on their relative copy numbers. Indeed, without explicit knowledge of $C$ (which depends, among other factors, on the number of cells involved in the experiment, and so is typically difficult to obtain), any absolute copy number profile consistent with the Hi-C map can be scaled by an integer and remains consistent.

2. *Absolute copy numbers only involve the products of mixture proportions and copy numbers, not either of them individually.* Since $\pi$ involves products of proportions and copy numbers, an increase in one can often be compensated by a decrease in the other without affecting even absolute values $\pi$ (i.e., this type of unidentifiability is independent of the scaling factor $C_0N$). More concretely, for any genome $G_0 \in \mathcal{G}$,

$$\pi = \sum_{G \in \mathcal{G}} f^G p^G = f^{G_0} p^{G_0} + \sum_{G \in \mathcal{G}_{\overline{0}}} f^G p^G = \left(1 - \sum_{G \in \mathcal{G}_{\overline{0}}} f^G\right) p^{G_0} + \sum_{G \in \mathcal{G}_{\overline{0}}} f^G \left(p^{G_0} + p^G - p^{G_0}\right)$$
$$= p^{G_0} + \sum_{G \in \mathcal{G}_{\overline{0}}} f^G \left(p^G - p^{G_0}\right) = p^{G_0} + \sum_{G \in \mathcal{G}_{\overline{0}}} \left(\frac{f^G}{K^G}\right) \cdot K^G \left(p^G - p^{G_0}\right),$$

where $\mathcal{G}_{\overline{0}} = \mathcal{G} \setminus \{G_0\}$ and $K^G \in \mathbb{N}$ are genome-specific constants. That is, even if the copy number profile for a single genome $G_0$ is completely known (in, e.g., most samples not derived from cell lines, it is reasonable to assume the presence of the purely diploid reference genome), the relative difference of all other genomes to $G_0$ are, in the most general setting, not determined by $\pi$. For the simplest case of two distinct cell populations, one purely diploid and the other with absolute copy number profile $p$, mixed at proportions $1 - f$ and $f$, respectively, the effective copy number profile reads

$$\pi_i = 2(1 - f) + fp_i = 2 + (p_i - 2)f.$$

That is, given the exact absolute copy number, it is only possible to infer the product $(p_i - 2)f$, and not either of them individually.

Given these fundamental limitations we must impose additional suitably restrictive, yet biologically plausible, assumptions. HiDENSEC does so by positing that:

(a) The most common effective copy number is known in advance; i.e., one has an estimate of

$$\pi_{\text{mode}} = \underset{\tau \in \bigcup_i \pi_i}{\arg\max} \# \{i : \pi_i = \tau\}.$$

Where $\# \{i : \pi_i = \tau\}$ is the number of bins with effective copy number $\tau$. This assumption allows rescaling of the $H_i$ to correct for the factor of $C_0N$. While we could use mean or median of $\pi$ for this purpose the mode of $\pi_{\text{mode}}$ is particularly appealing in that

1. unless the genomes are extraordinarily complex, $\pi_{\text{mode}}$ will be 2, and
2. $\# \{i : \pi_i = \pi_{\text{mode}}\}$ is often large, so that rescaling is effectively based on averaging many noisy observations, which in general outperforms estimation based on a single observation (as is done when only considering $\pi_1$).

(b) Copy number profiles are as close to purely diploid as is consistent with the data, in that $\max_{i,G} p_i^G$ is chosen as small as possible. For instance, consider the hypothetical example of $\mathcal{G} = \{G_0, G_1\}, p^{G_0} \equiv 2, \pi_i = 2 \cdot \mathbb{1}_{\{1,\ldots,i^*\}}(i) + 2.5 \cdot \mathbb{1}_{\{i^*+1,\ldots\}}(i)$, HiDENSEC will estimate

$f^{G_0} = 0.5, f^{G_1} = 0.5, p^{G_0} \equiv 2, p_i^{G_1} = 2 \cdot \mathbb{1}_{\{1,\ldots,i^*\}}(i) + 3 \cdot \mathbb{1}_{\{i^*+1,\ldots\}}(i)$ rather than, say, $f^{G_0} = 0.9, f^{G_1} = 0.1, p^{G_0} \equiv 2, p_i^{G_1} = 2 \cdot \mathbb{1}_{\{1,\ldots,i^*\}}(i) + 7 \cdot \mathbb{1}_{\{i^*+1,\ldots\}}(i)$, which is consistent with the principle of parsimony. Prior knowledge favouring a non-parsimonious solution can be explicitly fed into HiDENSEC as an optional argument if desired.

Although these assumptions narrow down the feasible solutions substantially, it can be shown that, in the most general setting, obtaining a solution is computationally intractable:

**Theorem.** *Identifying the smallest number of genomes $|\mathcal{G}|$ that explain a given noise-less effective copy number profile $\pi$ using mixture proportions bounded away from zero (e.g.,, $\min_{G\in\mathcal{G}} f^G \geq o(|\mathcal{G}|^{-1})$) and bounded absolute copy numbers (i.e., $\max_{G\in\mathcal{G}} \|p^G\|_\infty \leq B$ for some $B \in \mathbb{N}$) is, in general, at least as hard as the subset sum problem, and therefore NP-complete.*

Even though its proof does not immediately inform inference (and is therefore deferred to Section 5), this theorem suggests that any feasible inference procedure must be based on either

1. assumptions that are strong enough to render the subset sum problem efficiently solvable, yet are still biologically plausible, or

2. approximate inference which may work well for few genomes (e.g., $|\mathcal{G}|$ is small) but may become inaccurate as $|\mathcal{G}|$ grows large.

Given that empirically $|\mathcal{G}| = 2$ appears to typically explain the data well, with $|\mathcal{G}| > 3$ rarely being required (indeed, none of the cases described in the main text requires more than three genomes, HiDENSEC assumes a modest number of cancer genomes, and proceeds as follows:

1. Normalize the data $\{H_{ii}\}_i$ by $\hat{H}_{\text{mode}}/\pi_{\text{mode}}$, where $\hat{H}_{\text{mode}}$ is an estimate of the mode of $\{H_{ii}\}_i$. Data is typically abundant enough that obtaining $\hat{H}_{\text{mode}}$ through either a kernel density estimate or a simple histogram is sufficient. This normalized data is referred to as $\{\Pi_i\}_i$.

2. For a fixed window size $w$ (in the main text analysis $w = 100$), choice of $f$, and candidate copy number $p$, we define $\rho(f,p) = (1-f)2 + fp$ and

$$m_x(f,p) = \frac{1}{2w+1} \sum_{i=x-w}^{x+w} |\Pi_i - \rho(f,p)| \cdot \left[p \cdot \text{md}\left(\Pi_{[x-w,x+w]}\right)\right]^{-1},$$

where $\text{md}(X)$ is the median deviation of a set of numbers $X$, and the normalization involving it is motivated by the heteroskedasticity observed in contact-intensity counts (see **Supplementary Figure 12B**). For a choice of maximum copy number $p_{\max}$, compute a first estimated copy number profile $\hat{p}_x^1$ and associated mixture proportion $\hat{f}^1$ as the minimizers of $m_x(f,p)$ aggregated over the entire genome

$$\hat{f}^1 = \arg\min_f \sum_x \left[\min_{p\in[p_{\max}]} m_x(f,p)\right] \qquad \hat{p}_x^1 = \arg\min_{p\in[p_{\max}]} m_x(\hat{f}^1, p),$$

and estimate the corresponding effective copy number profile $\hat{\pi}^1$ as $\hat{\pi}^1 = \hat{f}^1 \hat{p}^1$. Estimation based on $m_x(f,p)$ in this manner exploits the strong spatial correlation present in $\Pi$, while otherwise remaining fully non-parametric.

3. Refine $\hat{\pi}_x^1$ by adjusting points of copy number changes, measuring their significance, and fine-tune $p_{\max}$ (see section below). Call this refined profile $\{\hat{\pi}_x^1\}_x$ as well.

4. If $\hat{\pi}^1 \equiv 2$, then return $\hat{\pi}^0 \equiv 2$ and $\hat{f}^0 = 0$.

5. Otherwise repeat steps 2-5 on the corrected effective copy number profile $\Pi^1 = \Pi - \hat{\pi}^1$ until $\hat{\pi}^K \equiv 0$, and then return $\hat{f}^{\mathcal{G}} = \bigcup_{k\in[K]} \hat{f}^k$, $\hat{p}^{\mathcal{G}} = \bigcup_{k\in[K]} \hat{p}^k$ and $\hat{\pi} = \sum_{k\in[K]} \hat{f}^k \hat{p}^k$.

In this way, HiDENSEC attempts to greedily explain the shape of $\Pi$ by subtracting the effect of an individual genome one at a time. It can be shown that this greedy procedure accurately recovers ground-truth $f^{\mathcal{G}}$ and $p^{\mathcal{G}}$ in the limit of noiseless data if the following conditions are met:

(a)
$$\left| \operatorname{supp} p^{G_k} \setminus \bigcup_{i=k+1}^{K} \operatorname{supp} p^{G_i} \right| \geq \left| \bigcup_{i=k+1}^{K} \operatorname{supp} p^{G_i} \right|,$$

for all $k \in [K]$, where $\operatorname{supp} p = \{i : p_i \neq 2\}$, and the $G_k$ are ordered such that $f^{G_1} \geq f^{G_2} \geq \cdots \geq f^{G_K}$.

(b)
$$2 \left\| \rho\left(f^{G_{k+1}}, ..., f^{G_K}, p^{G_{k+1}}, ..., p^{G_K}\right) \right\|_\infty \leq f^{G_k},$$

also for any $k \in [K]$, where by slight overloading of notation, $\rho\left(f_1, ..., f_r, p_1, ..., p_r\right) = 2\left(1 - \sum_{k=1}^r f_k\right) + \sum_{k=1}^r f_k p_k$.

(We note that if these conditions are not met, then the results of HiDENSEC may not be accurate.)

It is clear that these conditions tend to more easily be met if $K$ is small, while they become more restrictive as $K$ grows. Intuitively, they stipulate that genomes of more abundant cell populations should exhibit more substantial copy number changes than those of rare populations, and that mixture proportions be far away from uniformity; which—given the nature of logistic growth, and the fact that more abundant cell types likely had more time to evolve—appear biologically plausible. Indeed, these assumptions are satisfied in all samples analyzed in the main text (the majority of which carries $K = 1$ cell population in addition to the reference genome, in which case these conditions are trivially satisfied).

#### Refining effective copy number profiles

Due to the randomness inherent in chromatin folding and Hi-C experiments, the procedure described in step 2 above will occasionally detect copy number changes that are either imprecisely located or purely the result of stochastic fluctuation rather than biologically meaningful structure, and so it is desirable to correct for such misinference. HiDENSEC does so in various ways.

1. *Refining change points:* A site $x$ at which copy numbers change is characterized by the fact that $\mathbb{E}\Pi_{x-\delta} = \pi_{x-\delta} \neq \pi_{x+\delta} = \mathbb{E}\Pi_{x+\delta}$, for any sufficiently small $\delta$. If this gap between $\pi$ left

and right of $x$ is sufficiently large compared to the variance of the data, then it is reasonable to assume that

$$x = \underset{j \in \mathcal{N}(x)}{\arg\min}\, \mathbb{E}\,\mathrm{Var}\left[\Pi_\sigma \mid \mathbb{1}_{[-\infty, j-1]}(\sigma)\right], \tag{3}$$

where $\mathcal{N}(x)$ is a suitably small neighbourhood around $x$, and $\sigma \sim \mathrm{Uniform}\,(\mathcal{N}(x))$ is a uniform draw from $\mathcal{N}(x)$. That is, once likely change point candidates have been identified in step 2 above, their precise location can be refined by choosing suitable neighbourhoods $\mathcal{N}$ around them, and optimizing Eq. (3) accordingly (this essentially corresponds to fitting a depth-1 decision tree regressing sites in $\mathcal{N}$ against $\Pi$). The resulting refined change points are further adjusted or shifted to ensure that corresponding excursions (see below) do not cross chromosome boundaries.

2. *Interpreting change points:* The optimization procedure described above will return a refined choice of $x$ even if $\mathcal{N}$ does not undergo any copy number change, and so it is of interest to quantify the extent to which $x$ separates copy number levels. To do so, HiDENSEC assesses significance by performing 100 replicates of a permutation test on $\mathcal{N}$, and computing the $p$-value $p_{\mathcal{N}}(x)$ of $\mathrm{Var}\,[\Pi \mid \sigma_x]$ on the resulting empirical distribution. It should be noted that a priori it is unclear whether $p$-values calculated in such manner are well-calibrated even in the limit of large $\mathcal{N}$ (indeed, they should instead be formed based on the null distribution of counts around $\pi = 2$ conditional on $\hat{\pi} \neq 2$; which, however, is not accessible), but they do behave super-uniformly empirically (see **Supplementary Figure 12C**).

3. *Interpreting excursions:* An excursion $e$ of an effective copy number profile $\hat{\pi}$ is defined to be any tuple $e = (x_1, x_2, \hat{\pi}_{x_1+1})$ for two adjacent change points $x_1$ and $x_2$, for which $\hat{\pi}_{x_1+1} \neq 2$. $e$ is likely to be reflective of actual biological signal if $\max_{x \in \{x_1, x_2\}} p_{\mathcal{N}}(x)$ is small, if the length $x_2 - x_1$ of $e$ is large, and if the aggregate read counts on $[x_1, x_2]$ are broadly no more variable than expected for level $\hat{\pi}_{x_1+1}$ (if they are significantly more variable, then the change in effective copy number is prone to being merely a result of fluctuation). HiDENSEC thus assigns a significance to each excursion $e = (x_1, x_2, \pi)$ by incorporating the two $p$-values of both $x_1$ and $x_2$, one $p$-value associated with $x_2 - x_1$ based on a reference diploid genome, as well as one calculated from the median deviation of $\Pi$ on $e$ in relation to appropriately re-scaled diploid $\Pi$ values in the vicinity of $e$ (since read count fluctuations generally exhibit spatial dependence, with stochasticity increasing in smaller chromosomes, it is preferable to construct local empirical null distributions over global ones). Under $\mathcal{H}_0$, $p$-values computed in such a manner on a given set $\mathcal{E}$ of excursions behave broadly uniformly (see **Supplementary Figure 12C**).

4. *Model selection:* To assess whether $\hat{\pi}$ likely captures true somatic copy number alteration, or simply overfits to a noisy $\pi \equiv 2$ profile, various empirically well-performing checks are in place. More concretely, a $\hat{\pi}$ instance is declared overfitting (and whence adjusted to $\hat{\pi} \equiv 2$) if it clears any three of the following criteria:

    - *Inferred mixture proportion $\hat{f} < 0.15$.* The amplitude $A$ of a length-$\ell$ excursion that is purely due to stochastic fluctuations decays broadly as $O(e^{-A^2 \ell})$, and so detecting excursions consistent with large $f$ is unlikely under $\mathcal{H}_0$.

- $\text{md}\,\Pi \geq \hat{f}$. Small mixture proportions are only reliably attributable to biological signal if the fluctuations in $\Pi$ are of smaller order.
- *Number of excursions* $\geq 60$. Under $\mathcal{H}_0$, small-amplitude excursions are typically frequent.
- $|\{e = (x_1, x_2, \pi_e) \in \mathcal{E} : x_2 - x_1 \leq 200\}| \,/\, |\mathcal{E}| > 1/2$. Under $\mathcal{H}_0$, lengths of excursions decay exponentially, and so most observed excursions ought to be short.
- *The Benjamini-Hochberg threshold calculated on p-values computed in step 3 above calls less than* 10% *of* $\mathcal{E}$ *significant at* $\alpha = 0.25$. Under $\mathcal{H}_1$, $p$-values tend to be strongly significant.
- *Fluctuation strength is not monotonically increasing with copy number.* Under $\mathcal{H}_1$, larger copy numbers are associated with larger fluctuations.
- $\#\,\{x : \hat{\pi}_x = \pi_{mode}\}$ *makes up less than* $\varphi$% *of all* $x$, where $\varphi$ is by default set to 50, but can be adapted based on prior knowledge. Overfitting will lead to erroneous excursions away from $\pi_{\text{mode}}$.

5. *Utilizing off-diagonal information:* The presence or absence of off-diagonal signal at either boundary of an excursion $e$ provides further evidence for the biological significance of $e$. Thus, uncertainty quantification of off-diagonal intensities (discussed in the following section) is incorporated into the overall computation of a $p$-value associated with an excursion $e$.

Illustrations of effective copy number profiles called in this manner on the data sets analyzed in the main text are given in **Supplementary Figure 13**.

#### Confidence intervals for mixture proportions $\hat{f}^G$

There are primarily two sources of noise that contribute to uncertainty in the proportion estimates $\hat{f}^G$:

1. The stochasticity of read counts conditional on $h$; that is $\varepsilon$.
2. Shifts in the expected intensities $h$ themselves, due to, e.g., uncaptured covariates.

The former is a commonly encountered complication in statistical inference and can be addressed by classical non-parametric tools like the bootstrap (or versions thereof; e.g., the block or sieve bootstrap to account for the lack of independence and identical distributions in $\varepsilon$; it is also this lack of regularity in $\varepsilon$ that prevents exploiting more explicit tools based on central limit arguments or semi-parametric assumptions), while the latter is more delicate: It includes systematic biases in the data that may be unique to $\mathcal{G}$ (and therefore $h$) itself, and therefore can be difficult to estimate. For instance, for a (ground-truth) effective copy number profile $\pi$ featuring two excursions $e_1$ and $e_2$ at levels $\pi_{e_1} = 2 + \phi - \delta, \pi_{e_2} = 2 + \phi + \delta$ (for some $\phi \in [0, 1]$ and small $\delta > 0$), both of identical length, it is reasonable to either infer $|\hat{\mathcal{G}}| = 3, \hat{f}^1 = \phi, \hat{f}^2 = \delta$, or $|\hat{\mathcal{G}}| = 2, \hat{f} = \phi$ and attribute the shifts by $\delta$ to systematic biases that have not been captured by the covariate correction described above. HiDENSEC will decide between these two situations based on the fluctuations around these effective copy number values both inside and outside of these excursions, but if the latter is returned, then the difference of $\delta$ ought to be reflected in any uncertainty quantification of $\hat{f}^G$. To do so,

HiDENSEC performs the following bootstrap procedure, estimating confidence intervals for each $f^k$ in turn, and decoupling each excursion $e \in \mathcal{E}$:

1. For each $e \in \mathcal{E}$ with $e = (x_e, y_e, \hat{\pi}_e = 2\hat{f}^0 + \hat{f}^1\hat{p}^1)$ (that is, every excursions in which only the first non-reference genome in $\mathcal{G}$ is not diploid) of length $n_e = y_e - x_e$, resample $B_e$ Bootstrap replicates $\Pi_e^{b*}, b = 1, ..., B_e$ (where $B_e$ is determined below) of $\Pi$ on $e$, and compute local proportion estimates $(\hat{f_e^1})^{b*}$ as the medians of $|\Pi_e^{b*} - 2|/f$.

    - The resample sizes $B_e$ are chosen as the closest integer to $Bn_e/(\sum_{e'} n_{e'})$, where $B$ is chosen as large as computationally feasible (by default $B = 10^3$).

2. Remove outliers from $\bigcup_{e,b}(\hat{f_e^1})^{b*}$ by truncating past 3.5 median deviations, and return a 95% confidence interval around the resulting distribution's median.

3. Repeat steps 1 and 2 on each higher-order $\hat{\pi}^k$ that is not uniformly diploid, incorporating for each $e$ whose effective copy number estimate is contributed to by $f^1, ..., f^k$ the previously estimated uncertainties of $f^1, ..., f^{k-1}$.

The weighting scheme in step 1 is designed so as to attribute more importance to longer excursions, as these contribute more heavily towards estimating $\hat{f}^G$, with the trimming of step 2 encouraging erroneously called excursions to be excluded. Resorting to previously estimated uncertainties in $f^1, ..., f^{k-1}$ when estimating confidence intervals for $f^k$ in step 3 is necessary as fluctuations of $\sum_{i=1}^k \hat{f}_k\hat{p}_k$ inform fluctuations in $\hat{f}^k$ only when the fluctuations of $f^1, ..., f^{k-1}$ are known. Confidence intervals computed in this manner are likely to be conservative (though proving so requires further assumptions on the uncaptured covariates), since the bootstrapping design above effectively simulates inference of $\hat{f}^G$ on each excursion individually, while HiDENSEC estimates $\hat{f}^G$ using all excursions jointly. Nevertheless, **Supplementary Figure 3** illustrates that the resulting confidence intervals are reasonably small whenever appropriate.

### 4 Inference of large-scale structural variants

Large-scale structural variants typically result in off-diagonal intensities arranged in either of the six patterns given in **Supplementary Figure 14**, which in the following will be referred to as $\mathcal{P} = \mathcal{P}_1 \cup \mathcal{P}_2 = \{$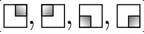$\} \cup \{$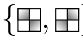$\}$. While events in $\mathcal{P}_1$ are most often associated with changes in the copy number profile, rearrangements falling into $\mathcal{P}_2$ typically are not, and so HiDENSEC treats their analysis separately. In particular, while HiDENSEC largely relies on its previously inferred copy number profiles $\hat{\pi}$ for detecting the former, the latter are called primarily based on their characteristic diagonal shape.

#### Detecting patterns in $\mathcal{P}_1$

Hi-C sub-matrices structured like the patterns in $\mathcal{P}_1$ can be found abundantly throughout the entire genome, and often correspond to intrinsic DNA geometry, compartment structure, or simply stochastic fluctuations inherent in the underlying biological and experimental processes. Moreover, in particularly noisy data or comparatively complex rearrangements, the area of largest intensity in any given $p \in \mathcal{P}_1$ may not be straightforward to identify, in which case all $p, q \in \mathcal{P}_1$ are approximately related to each other by a translation, and assigning one of them to a given empirical Hi-C sub-matrix may be under-determined. HiDENSEC addresses these two sources of uncertainty in two ways:

1. By default, HiDENSEC only reports off-diagonal events associated with excursions and corresponding copy number change points that have been evaluated as significant under the hypothesis testing scheme described in Sec. 3. Switching to non-default behavior and scanning arbitrary points along the genome is possible, but care should be taken in interpretation, as off-diagonal squares of enriched read counts may be confounded by above-mentioned biological and experimental hidden covariates. Reliably distinguishing signal due to noise from signal due to genomic rearrangement is often difficult even under manual detection by experts.

2. Since biological or experimental noise rarely result in individual $\mathcal{P}_1$ patterns in isolation, but rather display effects that tend to propagate horizontally, vertically, and locally along the Hi-C matrix (see, e.g., $\chi_6^p$ of Sample 1-II in **Figure 4**), each candidate Hi-C sub-matrix $H[\mathcal{J}, \mathcal{K}]$ that may potentially contain signal reflecting large-scale structural variant is evaluated in comparison to all sub-matrices obtained by translating $H[\mathcal{J}, \mathcal{K}]$ vertically (i.e., $\{H[\mathcal{J}, \mathcal{K}+y]\}_y$), horizontally (i.e., $\{H[\mathcal{J}+x, \mathcal{K}]\}_x$), and locally (i.e., $\{H[\mathcal{J}+x, \mathcal{K}+y]\}_{x,y=-w}^{w}$ for some window size $w$). Only if a suitable summary statistic of $H[\mathcal{J}, \mathcal{K}]$ (to be discussed below) appears sufficiently significant in comparison to the entire class of shifted sub-matrices, is $H[\mathcal{J}, \mathcal{K}]$ declared as containing evidence of genomic rearrangement events.

More concretely, HiDENSEC proceeds as follows.

1. For a given set of excursions $\mathcal{E}$ and associated $p$-values as determined in Sec. 3, select their most significant subset through Benjamini-Hochberg on a given significance threshold $\alpha$ (by default $\alpha = 0.05$). Call $\mathscr{C}$ the set of boundary points of the so selected candidate excursions.

2. For a choice of weight $w = (w_1, w_2)$ and off-diagonal point $x = (x_1, x_2)$, define the four quadrants
$$Q_w^{jk}(x) = \{(x_1 + m, x_2 + n) : m \in [0, jw_1], n \in [0, kw_2]\},$$
for $j, k \in \{\pm 1\}$, and denote by $H_w(x)$ the associated empirical distribution
$$H_w(x) = \sum_{j',k' \in \cup_{j,k} Q_w^{jk}(x)} H_{j'k'} \delta_{j'k'}$$
of read count locations. If $X \sim H_w(x), Y \sim \text{Uniform}\left(\bigcup_{j,k \in \{\pm 1\}} Q_w^{jk}(x)\right)$ are random variables distributed according to $H_w(x)$ and the uniform measure on $[x_1 - w_1, x_1 + w_1] \otimes [x_2 - w_2, x_2 + w_2]$, respectively, then HiDENSEC considers as test statistics
$$S_w^1(x) = \eta\left(\mathbb{E}\left[X \mid \mathcal{Q}_w(x)\right]\right) \qquad S_w^{2,\tau}(x) = \mathbb{E}\,\mathrm{Var}\left[\mathbb{E}\left(H(Y) \mid Y_\tau\right) \mid \mathcal{Q}_w^\tau(x)\right],$$
for $\tau \in \{1, 2\}$, where $\eta(Z)$ is the entropy of the random variable $Z$, $\mathcal{Q}_w = \left\{Q_w^{jk}(x)\right\}_{j,k \in \{\pm 1\}}$, $\mathcal{Q}_w^1(x) = \left\{\bigcup_{k \in \{\pm 1\}} Q_w^{jk}\right\}_{j \in \{\pm 1\}}$, and $\mathcal{Q}_w^2(x) = \left\{\bigcup_{j \in \{\pm 1\}} Q_w^{jk}\right\}_{k \in \{\pm 1\}}$. That is, while $S_w^1(x)$

essentially captures the extent to which read counts tend to accumulate in only one of the quadrants, $S_w^{2,\tau}$ measures whether read counts, projected onto the $X_\tau$ coordinate, exhibit evidence of copy numbers changing at $x$ (cf. Eq. (3) in Sec. 3).

3. For each pair of boundary points $\{x_1, x_2\} \in \binom{\mathscr{C}}{2}$ that fall into distinct chromosomes, refine its location by maximizing $S_w^1(x_1, x_2)$ locally through, e.g., coordinate ascent, and denote the resulting $\binom{|\mathscr{C}|}{2}$ off-diagonal indices by $\mathscr{C}$ as well.

4. For each $\{x_1, x_2\} \in \mathscr{C}$, compute $p$-values $p_{x_1}^1(x_2)$ and $p_{x_2}^1(x_1)$ from comparing $S_w^1(x, y)$ against the empirical distributions $\hat{S}_w^1(x_1) = \{S_w^1(x_1, y)\}_y$ and $\hat{S}_w^1(x_2) = \{S_w^1(y, x_2)\}_y$, where the index $y$ ranges over all genomic locations not part of the chromosome containing $x_1$ and $x_2$, respectively.

5. Compute $p$-values $p^{2,\tau}(x_1, x_2)$ by comparing $S_w^{2,\tau}(x_1, x_2)$ against the permuted random variable $\tilde{X} \sim (\sigma_1 \circ X_1, \sigma_2 \circ X_2)$, where $\sigma_k$ is drawn uniformly from the symmetric group on $[-w_k, w_k]$.

6. Under the null hypothesis of $S_w^1(x)$ following either $\hat{S}_w^1(x_1)$ or $\hat{S}_w^1(x_2)$, and $X_1 \perp\!\!\!\perp X_2$, $H(X_\tau) \perp\!\!\!\perp \mathscr{Q}_w^\tau(x)$, $m_w(x_1, x_2) = \min\{p_{x_1}^1(x_2), p_{x_2}^1(x_1\}$ is super-uniform, while $p^{2,\tau}(x_1, x_2)$ are uniformly distributed, with all three quantities independent of each other (note that the optimization in step (3) may affect these properties slightly, though as long as the refinement is kept sufficiently local, its impact appears empirically negligible; see **Supplementary Figure 13**). HiDENSEC thus ranks candidates in $\mathscr{C}$ based on $p$-values associated with $m_w(x_1, x_2) + \sum_\tau p^{2,\tau}(x_1, x_2)$, and declares a set $\mathscr{C}_+ \subset \mathscr{C}$ of significant off-diagonal contacts by means of the Benjamini-Hochberg procedure.

7. For each site $x$ identified in step (1), denote by $\mathscr{C}_x \subset \bigcup_{c \in \mathscr{C}_+} c$ the set of all its refinements partaking in a significant pair, and extract a single refinement by computing a combined on- and off-diagonal statistic akin to Eq. (3) on each element in $[\min \mathscr{C}_x, \max \mathscr{C}_x]$.

#### Detecting patterns in $\mathcal{P}_2$

Patterns in $\mathcal{P}_2$ are typically not tied to changes in copy number profiles, and thus require a more global search than what was necessary in the case of $\mathcal{P}_1$. However, their characteristic block-diagonal shape is rather more rigid; e.g., translational and within-block rotational symmetries do not apply in the same manner they did in $\mathcal{P}_1$, which HiDENSEC exploits for their detection. More explicitly, HiDENSEC proceeds as follows.

1. For each pair of chromosomes $\{\chi_a, \chi_b\}$, denote by $H[\chi_a, \chi_b]$ the Hi-C sub-matrix recording all contacts between $\chi_a$ and $\chi_b$.

2. For a fixed choice of $r$ (by default, $r = 50$), convolve $H[\chi_a, \chi_b]$ by $r^{-2} \mathbb{1}_{[r]} \otimes \mathbb{1}_{[r]}$, where $\mathbb{1}_{[r]} \in \mathbb{R}^r$ is the all-ones vector, and replace each entry $h_{ij}$ of the resulting smoothed matrix $\tilde{H}[\chi_a, \chi_b]$ by $\mathbb{1}_{h_{ij} > m}$, where $m$ is the median of non-zero values in $\tilde{H}[\chi_a, \chi_b]$. Interpret the so-constructed matrix $\overline{H}[\chi_a, \chi_b]$ as an encoding for a graph $\mathscr{G}[\chi_a, \chi_b]$, whose vertices $v$ are labeled $\{1, ..., |\chi_a|\} \times \{1, ..., |\chi_b|\}$ and whose every pair of vertices $v, w$ is connected by an edge if $\min_{s \in \{v,w\}} \left\{ \overline{H}[\chi_a, \chi_b]_s \right\} = 1$ and $\|v - w\|_\infty = 1$.

3. Fix a number $C$ of candidates to be considered per chromosome pair $\{\chi_a, \chi_b\}$, and identify the $C$ largest connected components $\mathscr{K}_1[\chi_a, \chi_b], ..., \mathscr{K}_C[\chi_a, \chi_b]$ (ordered in decreasing size) of $\mathscr{G}[\chi_a, \chi_b]$.

4. For each component $\mathscr{K}_j[\chi_a, \chi_b]$ identified in the step above, extract the vertex $v_j[\chi_a, \chi_b]$, such that $v_j[\chi_a, \chi_b] = \arg\max_v \tilde{H}[\chi_a, \chi_b](v)$, and let $\mathcal{V}[\chi_a, \chi_b] = \bigcup_{j\in[C]} v_j[\chi_a, \chi_b]$ be the collection of these vertices.

5. For a choice of window size $w$, sites $x, y$ and $Q_w^{jk}, \mathcal{Q}_w, \mathcal{Q}_w^{\tau}, H_w, X, Y$ as introduced in the previous section, define the statistics $T_w^{1,\sigma}(x, y), T_w^{2,\sigma}, T_w^{3,\sigma}, \sigma \in \{\pm 1\}$ as

$$\sigma T_w^{1,\sigma} = \mathbb{P}\left[(X, X') \in \left(Q_w^{11}(x, y), Q_w^{-1-1}(x, y)\right)\right] - \mathbb{P}\left[(X, X') \in \left(Q_w^{-1+1}(x, y), Q_w^{+1-1}(x, y)\right)\right]$$
$$T_w^{2,\sigma} = \mathbb{E}\left[H_w(Y) \mid \|Y - (x, y)\|_\infty \leq 3, Y \in Q_w^{1\sigma} \cup Q_w^{-1-\sigma}\right]$$
$$\sigma T_w^{3,\sigma} = \tau_{12}\tau_{21} - \tau_{11}\tau_{22},$$

where $X'$ is an *iid* copy of $X$, and $\tau_{kj}$ is given by

$$\tau_{kj} = \tau\left(m, \mathbb{E}\left[H_w(Y) \mid \|Y - (x, y)\|_\infty = m, Y \in Q_w^{kj}\right]\right)_{m\in[\rho]},$$

with $\tau(\cdot)$ being Kendall's $\tau$, and $\rho$ a pre-specified radius.

6. For each $v \in \bigcup_{a,b} \mathcal{V}[\chi_a, \chi_b]$, refine its location by locally maximizing first $H_w(v)$, and then $T_w^{1,\sigma}(v)$. Call these two collections of refined vertices $\mathcal{V}^\sigma, \sigma \in \{\pm 1\}$.

7. For each $v \in \mathcal{V}^\sigma$, compute $(T_w^{j,\sigma}(v))_{j\in\{1,2,3\}}$, and construct $p$-values $p_w^{1,\sigma}(v), p_w^{2,\sigma}(v)$ for every $v$ with $T_w^{3,\sigma}(v)$ larger than some threshold $t$ (by default, $t = 5$) as

$$p_w^{j,\sigma}(v) = \Phi_{\mu^{j,\sigma},\nu^{j,\sigma}}\left(T_w^{j,\sigma}(v)\right),$$

where $\Phi_{\mu,\nu}$ is the CDF of a Gaussian distribution with expectation $\mu$ and variance $\nu$, and

$$\mu^{j,\sigma} = |\mathcal{V}^\sigma|^{-1} \sum_{v\in\mathcal{V}^\sigma} T_w^{j,\sigma}(v) \qquad \nu^{j,\sigma} = |\mathcal{V}^\sigma|^{-1} \sum_{v\in\mathcal{V}^\sigma} (T_w^{j,\sigma}(v) - \mu^{j,\sigma})^2.$$

8. Under the null hypothesis of $(X \mid H_w(X))$ being uniformly distributed on $\mathcal{Q}_w$, $T_w^{1,\sigma}$ and $T_w^{2,\sigma}$ become Gaussian as $w$ and $\rho$ increase, and so $p_w^{j,\sigma}$ is approximately calibrated (see **Supplementary Figure 15C**). HiDENSEC then uses $\min\{p_w^{1,\sigma}, p_w^{2,\sigma}\}$ to select off-diagonal locations $v$ likely to exhibit patterns in $\mathcal{P}_2$.

**Supplementary Figure 15A,B** demonstrates the power and accuracy of the selection scheme described above, showcasing its strong calibration, high sensitivity, and precise localization: Manual expert inspection of all Hi-C matrices analysed in the main text yielded three distinct type-$\mathcal{P}_2$ fusion events, two of which are associated with mixture proportions of $\approx 10\%$. HiNT does not identify these two events as such (likely precisely due to their small associated proportions), but does declare the remaining third event as significant (alongside a similar number of false positives as discussed in the section above); however, returning a location estimate that differs from the actual signal by about $\|\hat{x}_{\text{HiNT}} - x\|_1 \approx 67$MB. In contrast, HiDENSEC correctly identifies all three—and only these three; i.e., at zero false-positive rate—events as such, with its location estimates coinciding precisely with those obtained from visual inspection.

#### Benchmarking

In order to more thoroughly assess the performance of HiDENSEC relative to HiNT outside the context of cell lines, the same benchmarking procedure as displayed in **Figure 3** was employed on all analyzed samples. As **Supplementary Figure 16** demonstrates, the relative improvement in top-$k$ recall remains as pronounced as, if not more so, in the setting of cell lines.

### 5 Proof of Theorem

**Theorem.** *Identifying the smallest number of genomes $|\mathcal{G}|$ that explain a given noise-less effective copy number profile $\pi$ using mixture proportions bounded away from zero (e.g., $\min_{G\in\mathcal{G}} f^G \geq o(|\mathcal{G}|^{-1})$) and bounded absolute copy numbers (i.e., $\max_{G\in\mathcal{G}} \|p^G\|_\infty \leq B$ for some $B \in \mathbb{N}$) is, in general, at least as hard as the subset sum problem, and therefore NP-complete.*

*Proof.* The proof proceeds by reducing the subset sum problem to two variants of it, one of which will be directly reducable to identifying $|\mathcal{G}|$. It begins by recalling the subset-sum problem in one of its most commonly stated form (here referred to as $\mathbf{SSP}_0$):

**Definition ($\mathbf{SSP_0}$).** *Given a set $S \subset \mathbb{Q}_+$ of $K$ non-negative rational numbers, and a target $T \in \mathbb{Q}$, decide whether there exists a subset $R \subset S$, so that $\sum_{r\in R} r = T$.*

$\mathbf{SSP}_0$ is well known to be NP-complete, and so any reduction of it to a new task $\mathbf{P}$ will render $\mathbf{P}$ NP-hard. The $\mathbf{P}$ of interest in the case here is the following:

**Definition ($\mathbf{Min}_{|\mathcal{G}|}$).** *Given a profile of effective copy numbers $\{\pi_i\}_i$, determine the smallest set $\mathcal{G}$, so that $\pi = \sum_{g\in\mathcal{G}} f^G p^G$ for some mixture proportions $f^G$ and absolute copy number profiles $p^G$, with* $\min_{G\in\mathcal{G}} f^G \geq g(|\mathcal{G}|) \in o(|\mathcal{G}|^{-1})$ *and* $\max_{G\in\mathcal{G}} \|p^G\|_\infty \leq B$ *for some $B \in \mathbb{N}$.*

It is clear that $\mathbf{Min}_{|\mathcal{G}|} \in \mathbf{NP}$, and so reducing $\mathbf{SSP}_0$ to $\mathbf{Min}_{|\mathcal{G}|}$ suffices to show that it is NP-complete. To do so, two intermediary reductions are needed:

$$\mathbf{SSP}_0 \leq \mathbf{SSP}_1 \leq \mathbf{SSP}_2 \leq \mathbf{Min}_{|\mathcal{G}|},$$

where $\mathbf{SSP}_1$ and $\mathbf{SSP}_2$ are defined to be

**Definition ($\mathbf{SSP}_1$).** *Given a set $S \subset \mathbb{Q} \cap [0,1]$ whose elements are linearly independent in the $\mathbb{Z}/B\mathbb{Z}$-module $\mathbb{Q}$, less than $2g(K)$, and sum to less than or $1$; and a target $T$, deciding if there exists a subset $R \subset S$ for which $\sum_{r\in R} r = T$ is NP-hard.*

**Definition ($\mathbf{SSP}_2$).** *Given a set $S$ as in $\mathbf{SSP}_1$, and a target $T$, deciding if there exists a subset $R \subset S$, and multiplicities $m \subset \mathbb{N}^S$ for which $\sum_{r\in R} m_r r = T$ is NP-hard.*

Indeed, if $\mathbf{SSP}_2$ is known to be NP-hard, then hardness of $\mathbf{Min}_{|\mathcal{G}|}$ follows:

**Lemma.**

$$\mathbf{SSP}_2 \leq \mathbf{Min}_{|\mathcal{G}|}.$$

*Proof of lemma.* Given an instance of $\mathbf{SSP}_2$, enumerate the elements of $S$ as $\{s_k\}_{k\in[K]}$, and construct an effective copy number profile consisting of $\pi_k = s_k$ as well as $\pi_{K+1} = T$. Due to the linear independence and boundedness assumptions on $S$, any $\mathcal{G}$ explaining such $\pi$ must be of size at least $K$ (with $f^{G_k} = s_k$ for $k \in [K]$), and will be of size $K+1$ if and only if $T = \sum_{k=1}^{K} f^{G_k} p^{G_k} = \sum_{k=1}^{K} s_k p^{G_k}$ for some $p_{K+1}^{\mathcal{G}} \in \mathbb{N}^K$. That is, if a set of genomes $\mathcal{G}$ of size $|\mathcal{G}| = K$ explains $\pi$, then setting $m_k = p_{K+1}^{G_k}$ solves $\mathbf{SSP}_2$; while otherwise no solution to $\mathbf{SSP}_2$ exists. ⊡

Thus it remains to show that $\mathbf{SSP}_0 \leq \mathbf{SSP}_2$.

**Lemma.**

$$\mathbf{SSP}_0 \leq \mathbf{SSP}_1.$$

*Proof of lemma.* Given an instance of $\mathbf{SSP}_0$,

1. find an invertible linear transformation $\tau(x) = ax + b$ such that $\tau(S)$ satisfies the boundedness assumptions of $\mathbf{SSP}_1$, and $b = b_0 + 10^{-e_0}$ for some $e_0$ much larger than any of the $e_k$ discussed below,

2. replace each $s_k$ by two new elements

$$s'_k = s_k^0 + 10^{-e_k} \qquad\qquad s''_k = s_k^1 - 10^{-e_k},$$

   where $s_k^i$ are positive, $s_k^0 + s_k^1 = \tau(s_k)$, and $S' = \cup_{k\in[K]}\{s'_k, s''_k\}$ is linearly independent in the $\mathbb{Z}/B\mathbb{Z}$-module $\mathbb{Q}$, and $e_k \in \mathbb{N}$ are exponents larger than the maximum of $\mathcal{F}_{10}(aT + Kb)$ and $\max_{k\in[K],i\in\{0,1\}} \mathcal{F}_{10}(s_k^i)$ (where $\mathcal{F}_{10}(x)$ is the largest index—counting from the left—at which the base-10 expansion of $x$ is non-zero), distinct from each other; i.e., $e_k \neq e_\ell$ if $k \neq \ell$, and chosen so as to not violate any boundedness assumptions.

Then $(S', aT + kb))_{k\in[K]}$ are all valid instances for $\mathbf{SSP}_1$, and any solution must either select both or neither of $s'_k$ and $s''_k$. If one of these instances, say the $k_*^{\text{th}}$, accepts on a subset of indices $R$, then $|R| = k_*$ due to the choice of $e_0$, and since

$$\sum_{r\in R} as_r + b = k_* b + a \sum_{r\in R} s_r = aT + k_* b,$$

it must be true that $\sum_{r\in R} s_r = T$, and so $R$ too provides a positive answer to $(S, T)$. Conversely, a solution $R$ to $(S, T)$ will provide a solution to $(S', aT + |R|b)$, and so the lemma is proved. ⊡

**Lemma.**

$$\mathbf{SSP}_1 \leq \mathbf{SSP}_2.$$

*Proof of lemma.* A similar proof idea as in the lemma just proved works here as well: Each element $s_k \in S$ is replaced by two elements that indicate whether $s_k$ is used once or not at all in the following manner.

1. Choose $(e_k)_{k\in[K]}$, so that $e_k \geq B' + \max_{x\in S\cup\{T\}} \mathcal{F}_{10}(x)$, and such that $|e_k - e_\ell| \geq B'$ for some $B' > B$.

2. Replace each element $s_k$ with two elements

$$s'_k = 10^{-e_k} \qquad\qquad s''_k = s_k + 10^{-e_k}.$$

3. Define $S' = \cup_{s \in S} \{s'_k, s''_k\}$, $T' = T + \sum_{k \in [K]} 10^{-e_k}$, and query $\mathbf{SSP}_2$ on the instance $(S', T')$.

If $\mathbf{SSP}_2$ returns a solution $R'$ to this instance, then $R = \{k : s''_k \in R'\}$ provides a solution of indices to $\mathbf{SSP}_1$ on $(S, T)$. The converse direction is clear. ⊡

Chaining together these individual lemmas yields the theorem as desired. □

Although this proof may appear contrived on first glance, it in fact describes the very difficulty HiDENSEC must deal with: Given various levels $\pi_1, ..., \pi_K$ of $\pi$, can a new level $\pi_{K+1}$ be explained by the same genomes that explain $\pi_1$ through $\pi_K$ or is the introduction of a new one necessary? The proof shows that even when a set of genomes explaining $\pi_1, ..., \pi_K$ is known, answering this question in general is intractable—therefore, in practice, where the genomes explaining $\pi_1$ through $\pi_K$ are not known and must be estimated themselves, this must be true too.

### 6 Supplementary Figures

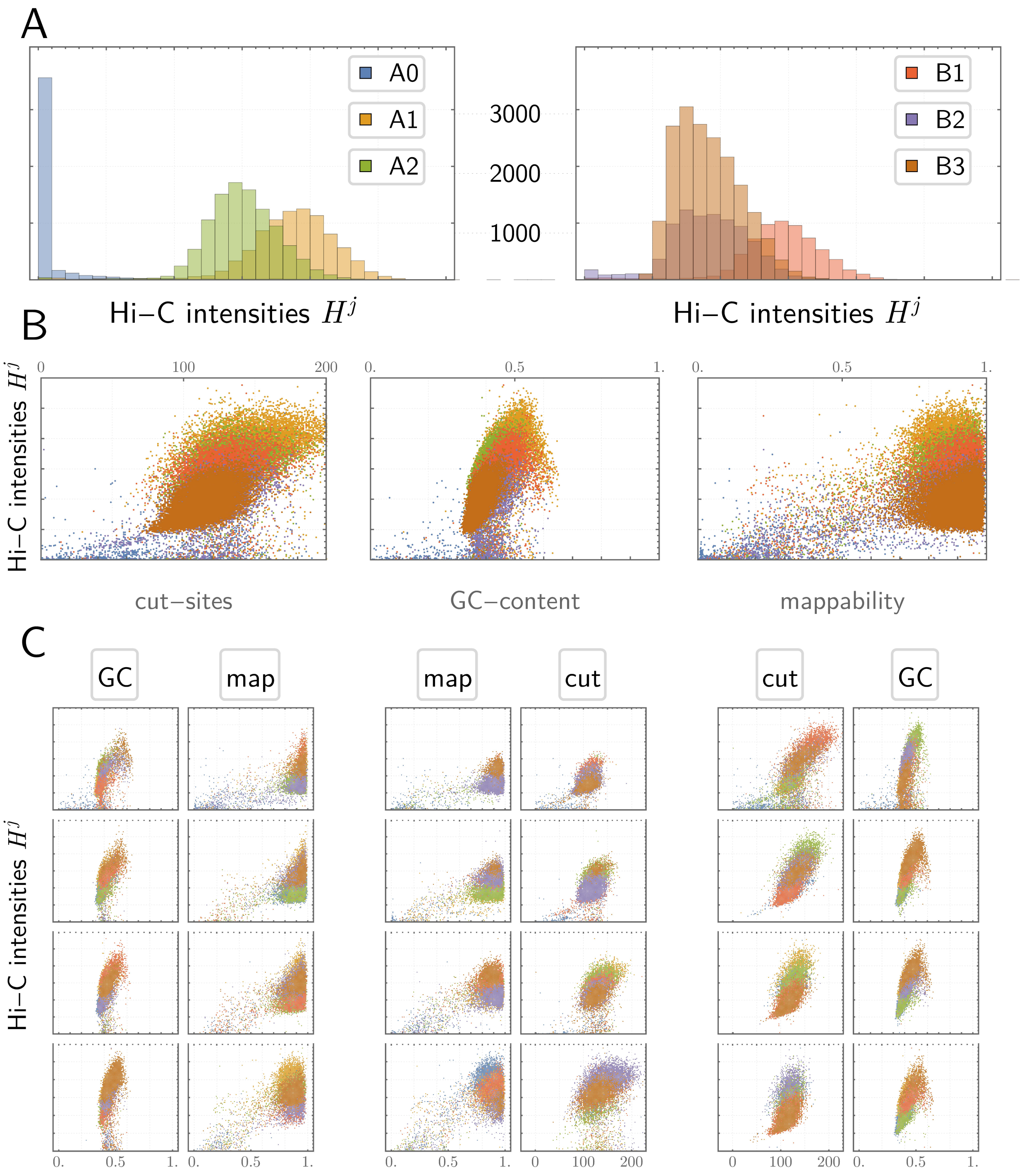


**Supplementary Figure 1: Covariate dependence of $H^j = \sum_k H_{jk}$ in GM12878 *in situ* Hi-C data**. The impact of the four covariates compartment structure, GC-content, number of cut-sites and mappability on row sums of Hi-C intensity matrices is displayed. **A**: $H^j$ conditioned on compartment structure, **B**: $H^j$ as a function of remaining three covariates; points are coloured by compartment, **C**: $H^j$ conditioned on quartiles of the corresponding column statistic in B, as a function of the two remaining covariates.

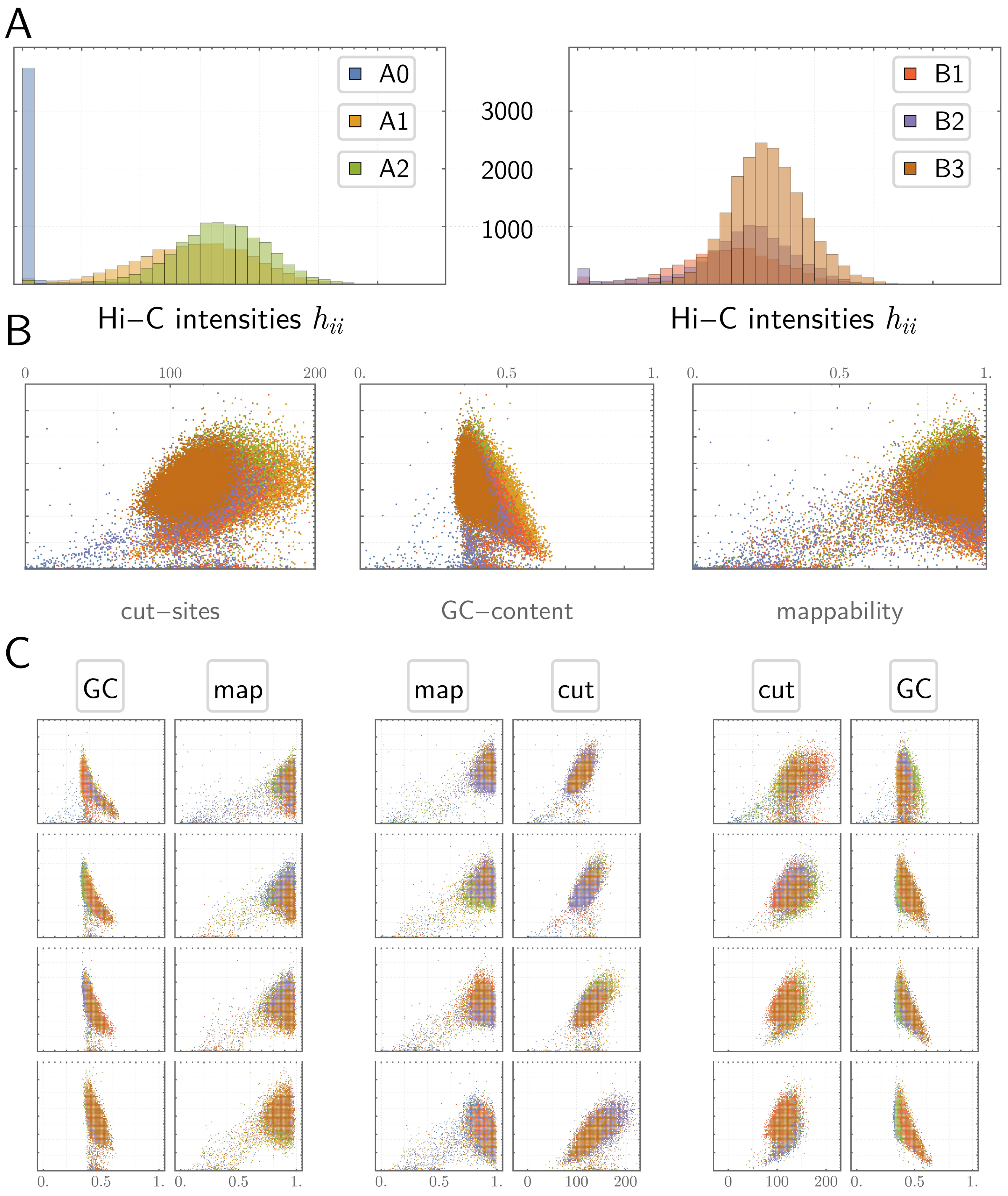


**Supplementary Figure 2: Covariate dependence of $H^j = \sum_k H_{jk}$ in Sample 1-I *in vivo* Fix-C data**. Plots are as described in **Supplementary Figure 1**.

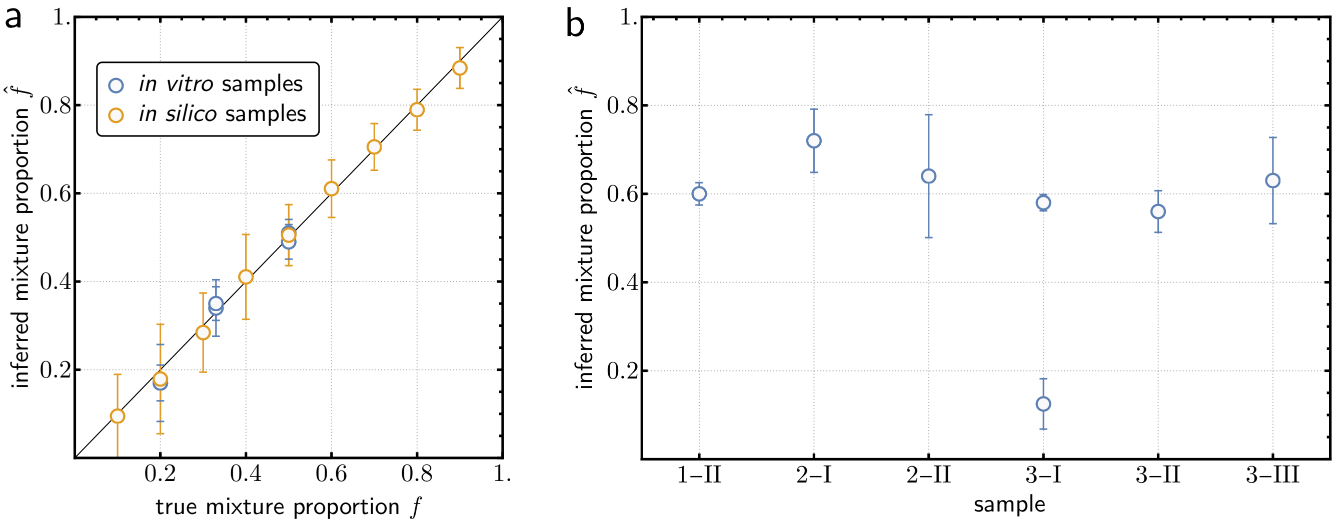


**Supplementary Figure 3: Predictions of tumor purity correlate well with known tumor purity.** (**a**) The x-axis represents true tumor purity for in vitro and in silico samples; while the y-axis represents the HiDENSEC inferred tumor purities. There is high concordance between the two and the error bars represent 95% confidence intervals for the HiDENSEC inferred tumor purity. (**b**) The x-axis represents different samples used in the analysis (1-II represents Sample 1-II, for instance). The y-axis represents the inferred tumor purity (the fraction of cells that are cancerous). The error bars represent 95% confidence intervals for the inferred tumor purity.

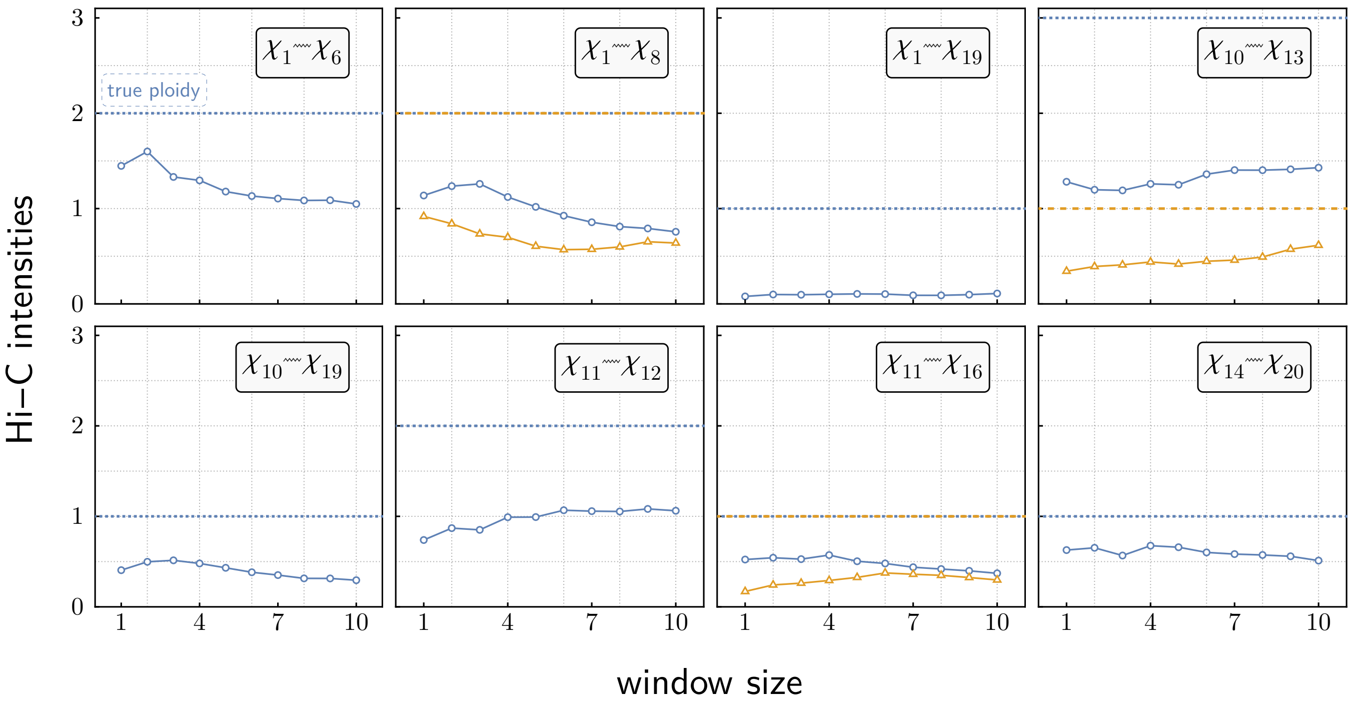


**Supplementary Figure 4: Off-diagonal intensities of large-scale chromosomal rearrangements aren't reliable estimators of absolute copy number.** Each panel represents the Hi-C intensities corresponding to a particular off-diagonal event in the HCC1187C cell line (for instance the top-left panel represents a translocation between chromosome 1 and chromosome 6). The horizontal axis represents a measure of how large a window around the translocation (which manifests itself as an off-diagonal event on the Hi-C map) was considered to compute the Hi-C intensity. The Y-axis represents the resulting Hi-C intensity. The true ploidies of inter-chromosomal translocations are denoted by the horizontal dotted lines while the colored curves represent the measured Hi-C intensities. For balanced translocations there are two colored lines corresponding to the two fusion events. The fact that the ratio of the true ploidy represented by the horizontal dotted lines and the colored curves is not consistent across the various large-scale chromosomal rearrangements within the same sample, suggests that the off-diagonal intensities are confounded by covariates in addition to absolute copy number and hence HiDENSEC does not use off-diagonal Hi-C intensities to infer absolute copy numbers.

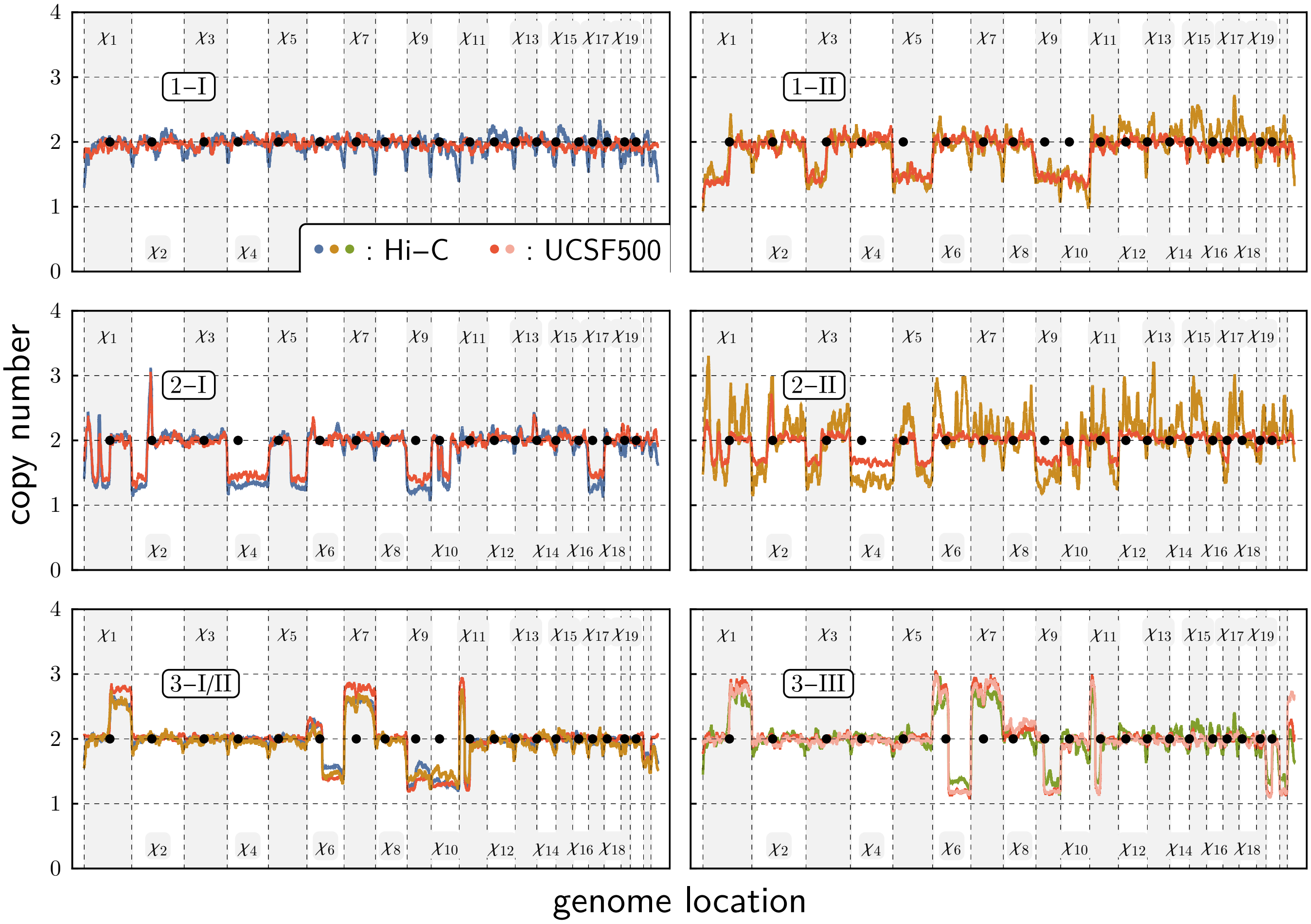


**Supplementary Figure 5: HiDENSEC absolute copy number predictions correlate well with genome-wide copy numbers inferred from next-generation sequencing.** Each panel represents HiDENSEC absolute copy number predictions for a particular sample (whose identity is indicated inside each panel; e.g., 1-II in the top-right represents Sample 1 - II), compared to UCSF500 or Exome sequencing based (relative) copy number calls. Copy number profiles inferred by HiDENSEC from Hi-C data use color codes consistent with the main text (that is, blue curves correspond to Samples 1 - I, 2 - I, 3 - I, beige curves to Samples 1 - II, 2 - II, 3 - II, and the green curve depicts Sample 3 - III), while red and pink profiles represent relative copy numbers (transformed by $x \to 2 \times 2^x$) from CNVkit using UCSF500 or Exome sequencing. For Sample 2 - I, the discordance between the levels of the HiDENSEC absolute copy numbers and the CNVkit scaled relative copy numbers inferred using UCSF500 data is likely due to differences in samples for the Hi-C data and for the UCSF500 data, since the UCSF500 based tumor purity lies outside the 95% confidence intervals of the HiDENSEC inferred tumor purity (**Supplementary Figure 3**, **Supplementary Table 4**). For Sample 3 - III, two UCSF500 based curves are displayed, as UCSF500 data from two different metastases corresponding to this sample exists; both of these are concordant with the HiDENSEC absolute copy number inferred using Hi-C data from the metastasis sample, Sample 3 - III.

a

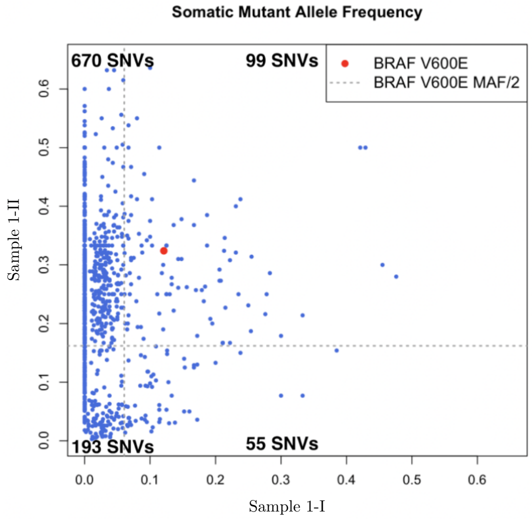


b

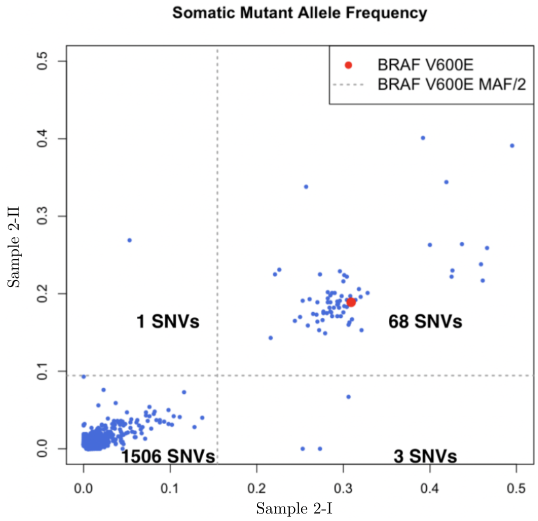


**Supplementary Figure 6: Somatic mutant allele frequencies for Patient 1 and Patient 2, based on somatic variant calls derived from Exome sequencing data and UCSF500 data, respectively. (a)** Somatic variant calls were obtained using Mutect2 from exome sequencing data in samples derived from Patient 1 and **(b)** UCSF500 data in samples derived from Patient 2. These somatic variant calls were then filtered out for false positives and the intersection of mutations observed in both samples within a patient was considered. The x-axis and the y-axis in each of the two panels represents somatic mutant allele frequencies. Each individual data point is a particular somatic variant call, from among the intersection of filtered somatic variant calls within the two samples of a patient. The red point denotes the well-known BRAF V600E somatic variant while the dashed lines are drawn at exactly half of its mutant allele frequencies. The resulting quadrants are intended to denote somatic variants common to both samples (corresponding to the top-right quadrant), somatic variants present in only one of the samples (corresponding to the top-left and the bottom-right quadrants), and somatic variants which are likely false positives (bottom-left quadrant).

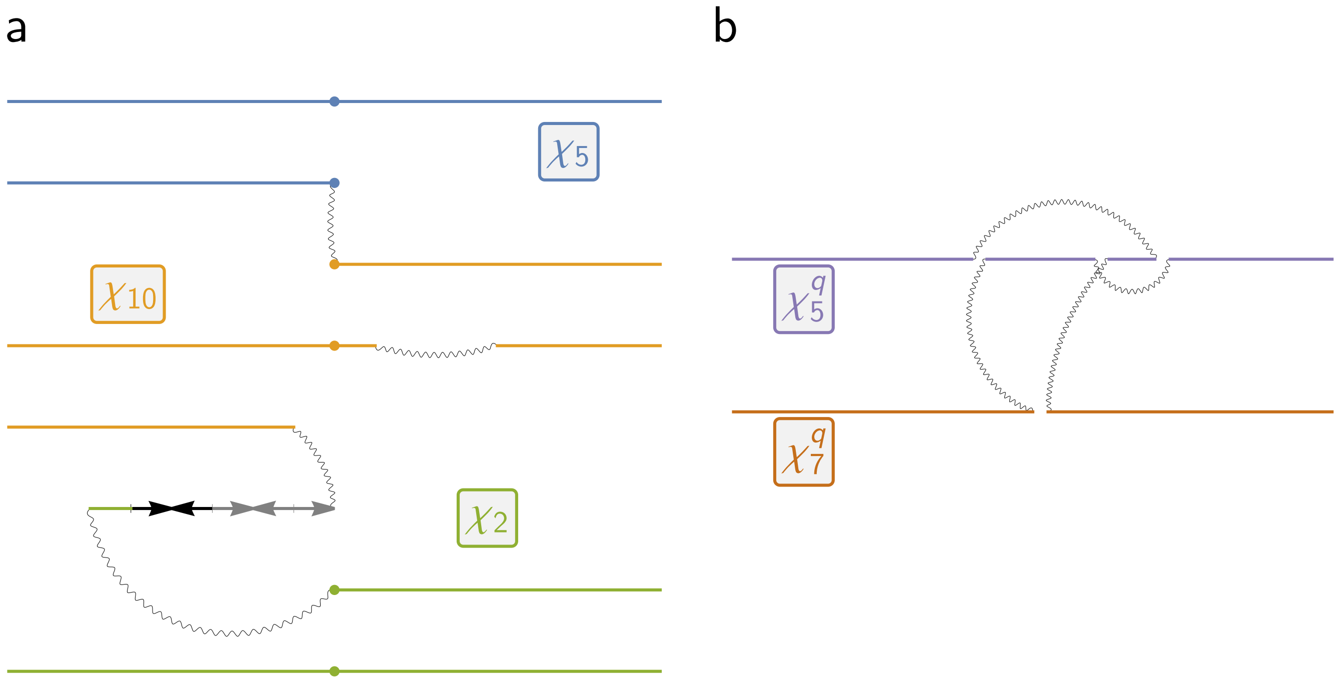


**Supplementary Figure 7: Schematics demonstrating two particularly complex structural variants.** The translocations involving chromosome 2, 5 and 10 in both samples of Patient 2 (Sample 2 - I and Sample 2 - II) and the inter-chromosomal translocation between chromosome 5 and 7 in Sample - I have been depicted here. In order to infer the contacts constituting these complex structural variants, the Hi-C maps of the relevant chromosomes were carefully analyzed in conjunction with the HiDENSEC inferred copy numbers, as shown in **Supplementary Figure 9, Supplementary Figure 10**. Arrows indicate duplications and inversions of genomic segments of chromosome 2.

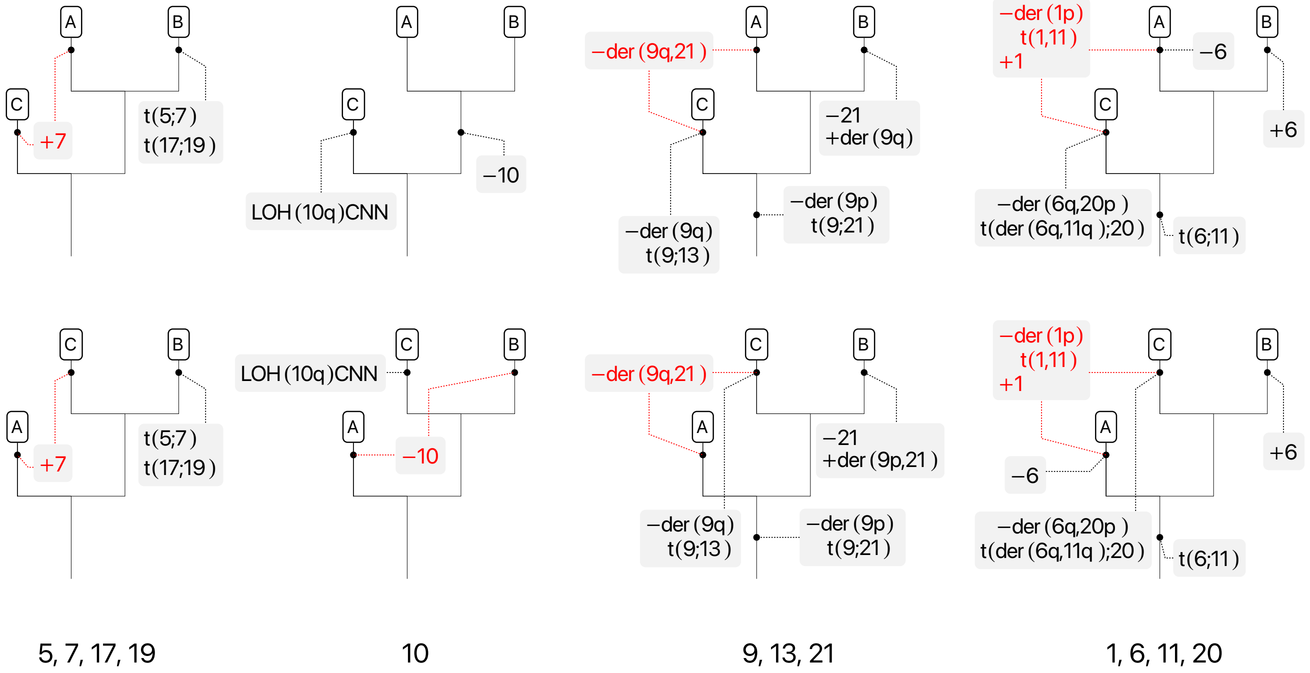


**Supplementary Figure 8: Alternatives to the phylogenetic relationship between the three cell types in Patient 3 .** Two alternative phylogenies that are consistent with the large-scale chromosomal rearrangements in Patient 3. The phylogeny in **Figure 7b** was chosen over these two alternatives following the principle of parsimony since the number of convergent events, denoted in red, are higher in these two phylogenies, than the one described in **Figure 7b**.

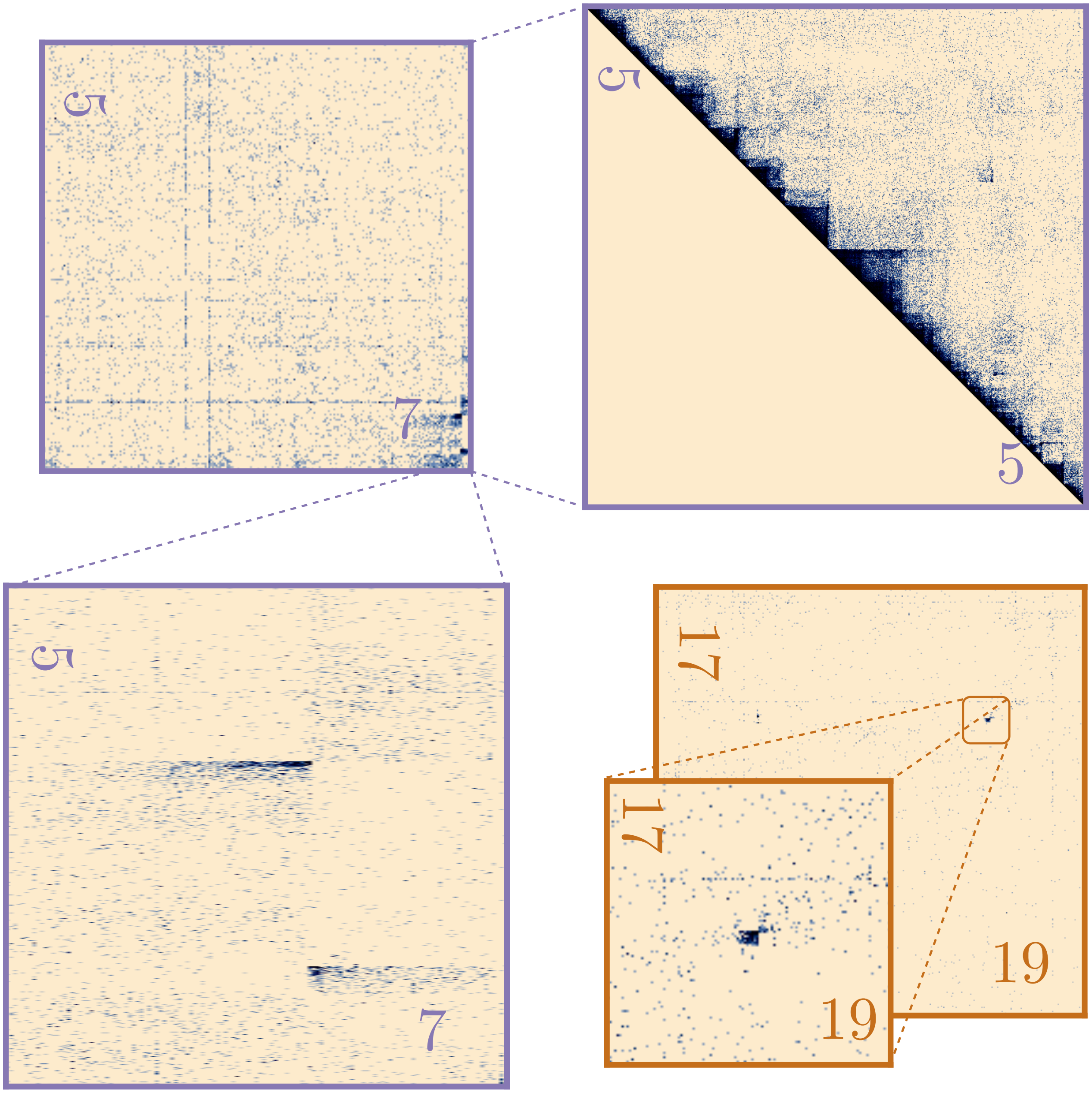


**Supplementary Figure 9: Zoomed-in Hi-C maps of two translocations observed in Sample 3 - I.** Sample 3 - I contains two structural variants that are relatively smaller in size than chromosome arms. The first of which is a complex structural variant between chromosome 5 and chromosome 7, a schematic of which is depicted in **Supplementary Figure 7b**. The second structural variant is a short balanced translocation between chromosome 17 and chromosome 19, depicted in the bottom right panel and its inset.

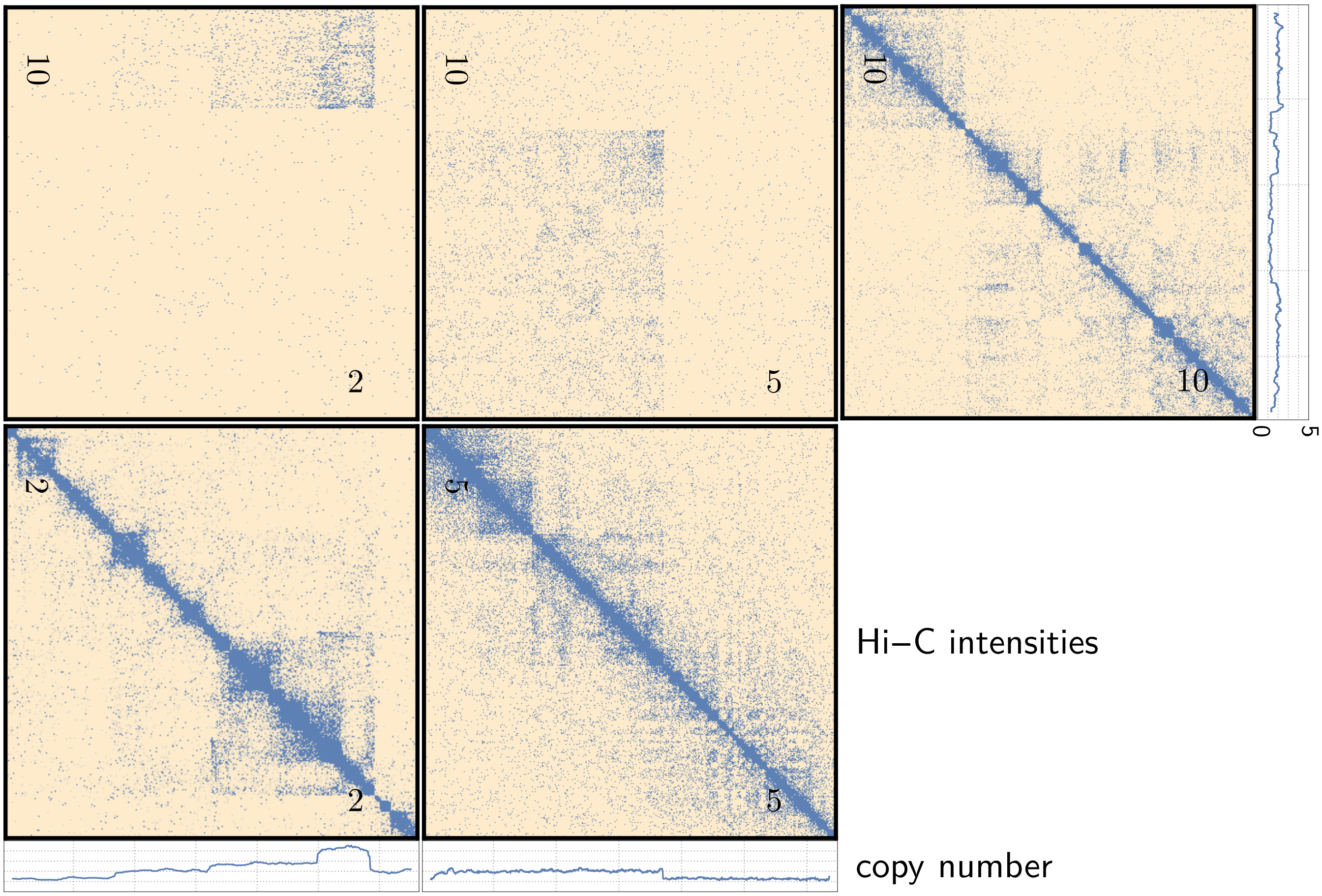


**Supplementary Figure 10: Hi-C contact maps used to infer the inter-chromosomal translocations involving chromosomes 2, 5 and 10 in Patient 2.** This figure shows the zoomed in Hi-C maps for chromosome pairs involved in the complex structural event depicted in **Supplementary Figure 7a**. The three line plots adjacent to the Hi-C maps represent HiDENSEC inferred copy numbers for the three chromosomes, which were used to determine the contacts constituting this complex structural variant.

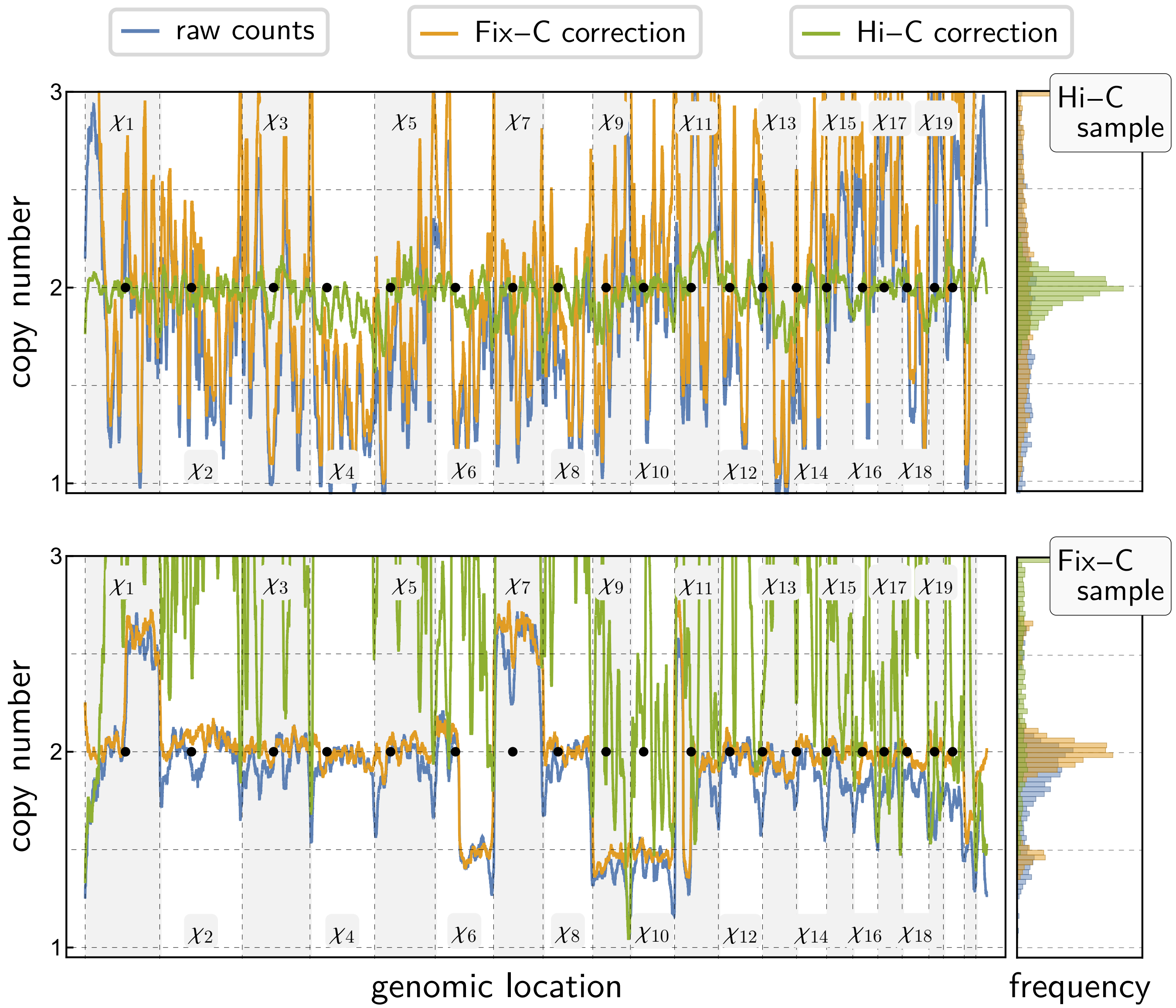


**Supplementary Figure 11: Covariate correction is protocol-dependent.** A Hi-C sample of the reference genome as well as the Fix-C Sample 3-II illustrate the necessity for both covariate correction in general, as well as its protocol-specific nature.

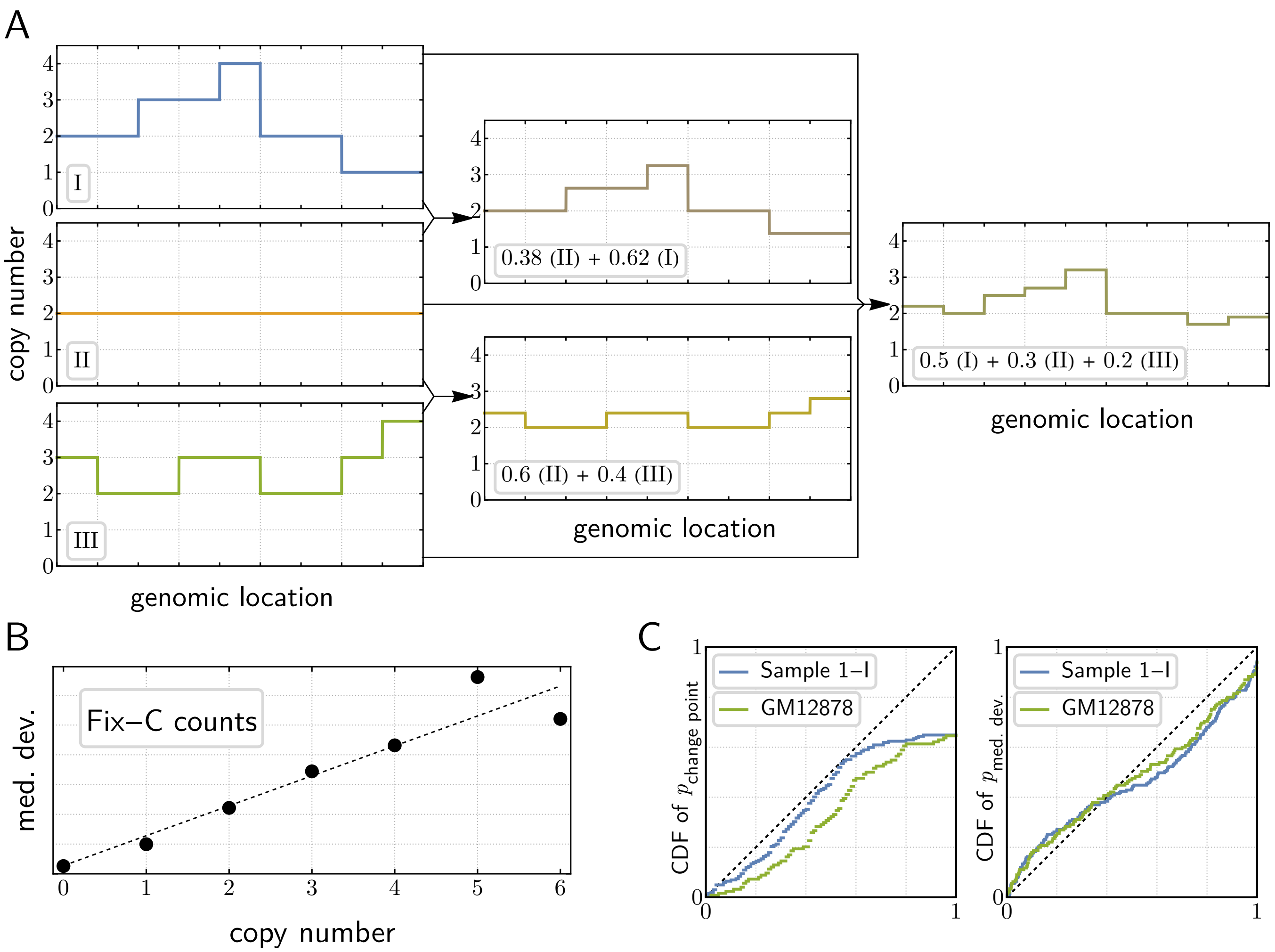


**Supplementary Figure 12: Generative model of signal & noise.** Observed on-diagonal contact intensities are modeled as an underlying effective copy number profile comprised of a convex combination of individual, cell-population-specific absolute copy number profiles (**A**), which is perturbed by heteroskedastic noise (**B**) and scaled by a generally unknown constant $C_0N$. Under $\mathcal{H}_0$ of $\pi \equiv 2$, $p$-values associated with HiDENSEC 's test statistics behave super-uniformly or close to uniform (**C**).

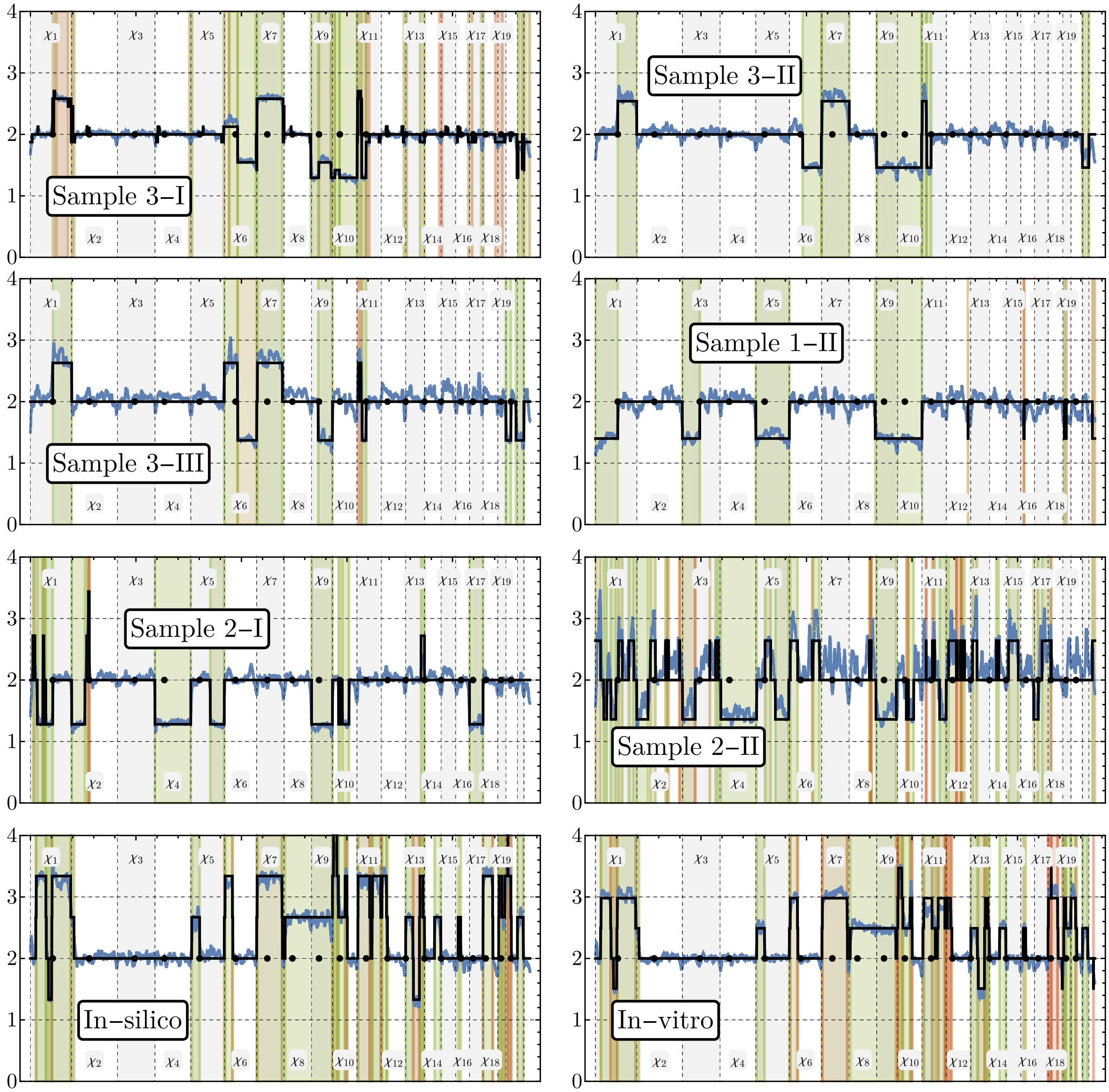


**Supplementary Figure 13: Inference of effective copy number profiles & interpretation of excursions.** Inferred $\hat{\pi}$ (solid black line) for each of the non-diploid samples discussed in the main text as well as for in-silico and in-vitro mixtures (with $f = 0.7$ and $f = 0.5$, respectively) are shown against $\Pi$ (blue line). Each excursion $e$ is associated with a $p$-value reflecting its biological significance, with greener colors mirroring higher significance.

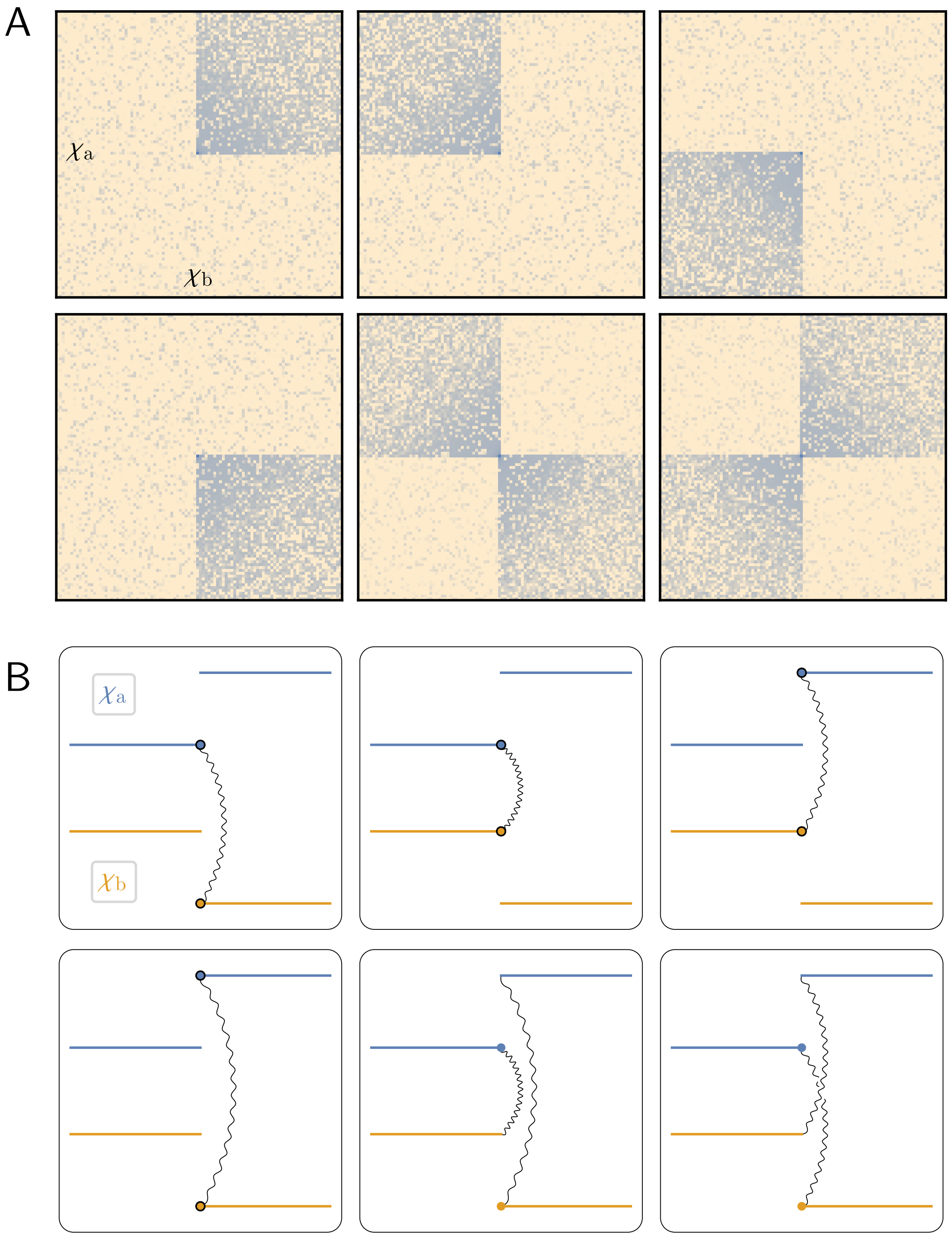


**Supplementary Figure 14: Hi-C intensity patterns and associated large-scale structural variants.** HiDENSEC detects Hi-C sub-matrices of six distinct patterns (**A**) associated with six types of large-scale structural variants (**B**) (note: non-fusing segments may interact with chromosomes other than $\chi_a$ and $\chi_b$ or be deleted without qualitatively affecting the local Hi-C patterns of (A)).

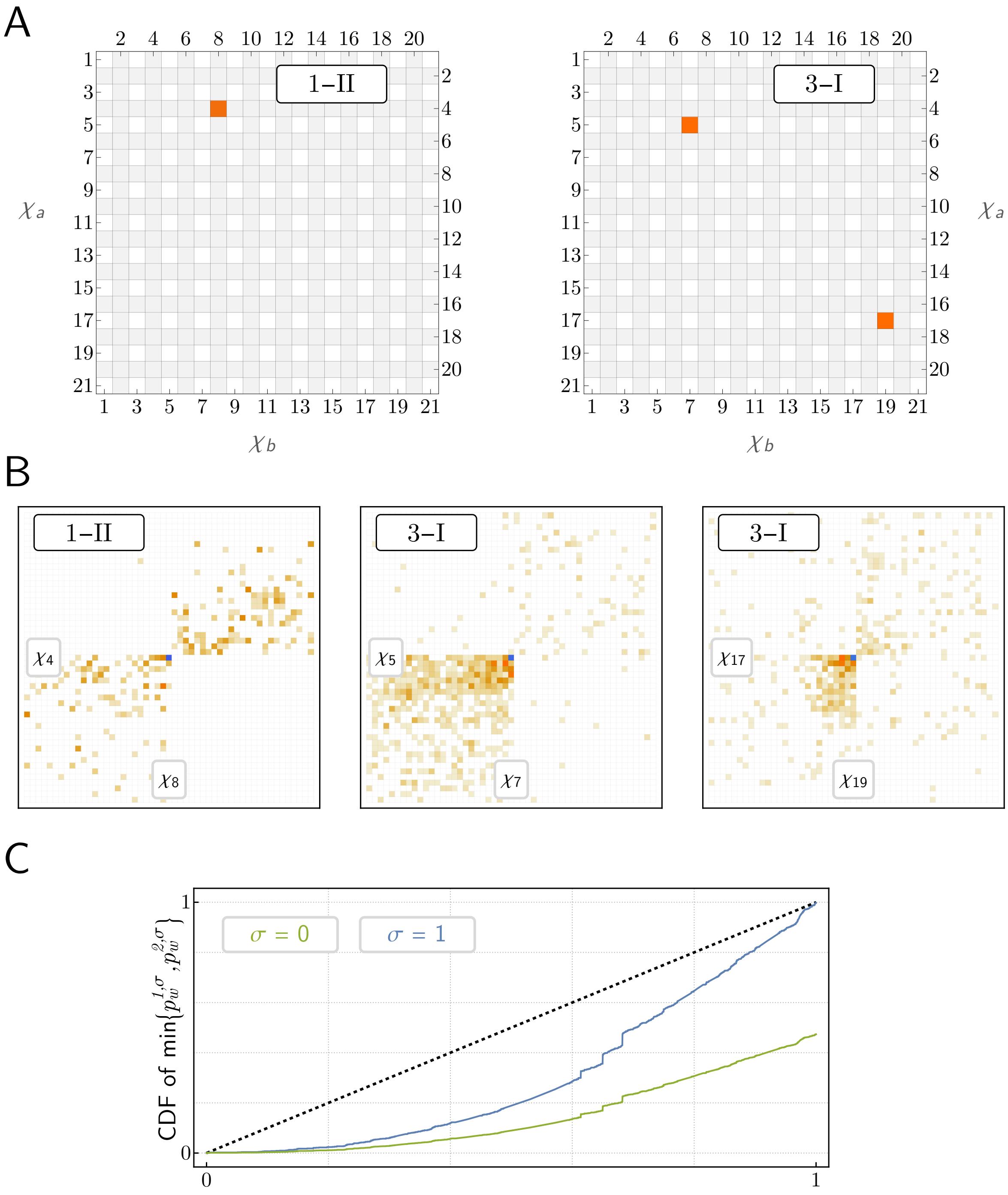


**Supplementary Figure 15: HiDENSEC reliably detects off-diagonal exchange patterns.** In those samples that do contain patterns in $\mathcal{P}_2$, HiDENSEC correctly recovers them at zero false-positive rate (**A**), and identifies their precise locations accurately (**B**, blue highlights indicate fusion sites inferred by HiDENSEC). Calibration of HiDENSEC is primarily a result of computed $p$-values behaving super-uniformly (**C**, empirical distributions based on all samples analyzed in the main text). Of the three events, HiNT only detected the $\chi_4 \sim \chi_8$ fusion, locating it, however, $\approx$ 42 and $\approx$ 25MB away from the true signal on $\chi_4$ and $\chi_8$, respectively.

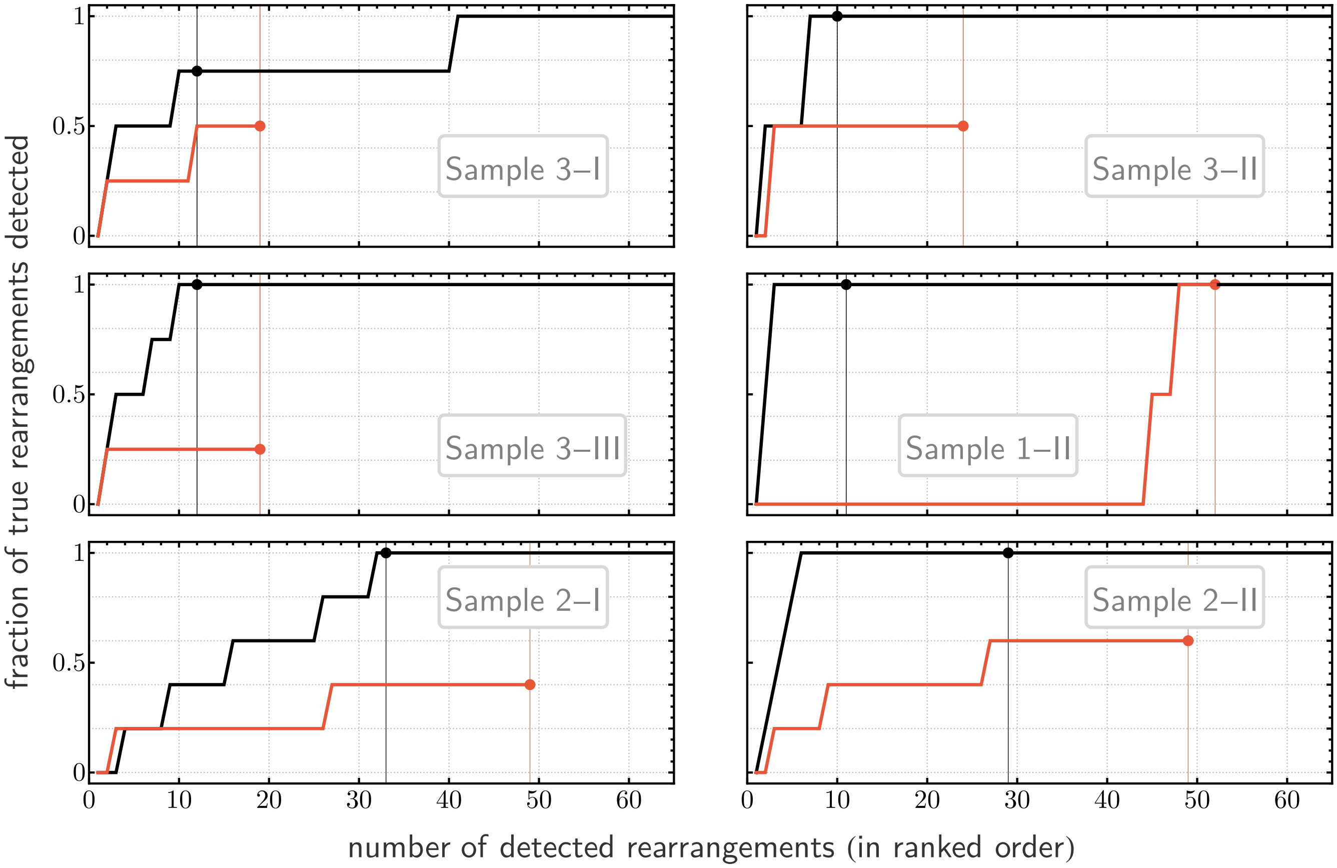


**Supplementary Figure 16:** Comparison of HiDENSEC's (black) and HiNT's (red) top-$k$ recall on the samples analyzed in the main text. As in the corresponding main figure, filled regions indicate rearrangements deemed significant by either method.
